# Data-driven detection of latent atrophy factors related to phenotypical variants of posterior cortical atrophy

**DOI:** 10.1101/679225

**Authors:** Colin Groot, B.T. Thomas Yeo, Jacob W Vogel, Xiuming Zhang, Nanbo Sun, Elizabeth C. Mormino, Yolande A.L. Pijnenburg, Bruce L. Miller, Howard J. Rosen, Renaud La Joie, Frederik Barkhof, Philip Scheltens, Wiesje M van der Flier, Gil D. Rabinovici, Rik Ossenkoppele

**Affiliations:** Alzheimer Center Amsterdam, Department of Neurology, Amsterdam Neuroscience, Vrije Universiteit Amsterdam, Amsterdam UMC, Amsterdam, The Netherlands; Department of Electrical and Computer Engineering, Clinical Imaging Research Centre, N1. Institute for Health and Memory Networks Program, National University of Singapore, Singapore, Singapore; Montreal Neurological Institute, McGill University, Montreal, QC, Canada; Computer Science and Artificial Intelligence Laboratory, Massachusetts Institute of Technology, Cambridge, MA, USA; Department of Neurology and Neurological Sciences, Stanford University, Stanford, CA, USA; Department of Radiology and Nuclear Medicine, Amsterdam Neuroscience, Vrije Universiteit Amsterdam, Amsterdam UMC, Amsterdam, The Netherlands; Institutes of Neurology & Healthcare Engineering, University College London, London, United Kingdom; Department of Epidemiology and Biostatistics, Amsterdam Neuroscience, Vrije Universiteit Amsterdam, Amsterdam UMC, Amsterdam, The Netherlands; Departments of Neurology, Radiology and Biomedical Imaging, University of California, San Francisco, San Francisco, CA, USA; Clinical Memory Research Unit, Lund University, Lund, Sweden

## Abstract

Posterior cortical atrophy is a clinical-radiological syndrome characterized by visual processing deficits and atrophy in posterior parts of the brain, most often caused by Alzheimer’s disease pathology. Recent consensus criteria describe four distinct phenotypical variants of posterior cortical atrophy defined by clinical and radiological features; i) object perception/occipitotemporal (ventral), ii) space perception/temporoparietal (dorsal), iii) non-visual/dominant parietal and iv) primary visual (caudal). We employed a data-driven approach to identify atrophy factors related to these proposed variants in a multi-center cohort of 119 individuals with posterior cortical atrophy (age: 64 SD 7, 38% male, MMSE: 21 SD 5, 71% amyloid-β positive, 29% amyloid-β status unknown). A Bayesian modelling framework based on latent Dirichlet allocation was used to compute four latent atrophy factors in accordance with the four proposed variants. The model uses standardized gray matter density images as input (adjusted for age, sex, intracranial volume, field strength and whole-brain gray matter volume) and provides voxelwise probabilistic maps for all atrophy factors, allowing every individual to express each factor to a degree without *a priori* classification. The model revealed four distinct yet partially overlapping atrophy factors; right-dorsal, right-ventral, left-ventral, and limbic. Individual participant profiles revealed that the vast majority of participants expressed multiple factors, rather than predominantly expressing a single factor. To assess the relationship between atrophy factors and cognition, neuropsychological test scores covering four posterior cortical atrophy-specific cognitive domains were assessed (object perception, space perception, non-visual parietal functions and primary visual processing) and we used general linear models to examine the association between atrophy factor expression and cognition. We found that object perception and primary visual processing were associated with atrophy that predominantly reflects the right-ventral factor. Furthermore, space perception was associated with atrophy that predominantly represents the right-ventral and right-dorsal factors. Similar to the atrophy factors, most participants had mixed clinical profiles with impairments across multiple domains. However, when selecting four participants with an isolated impairment, we observed atrophy patterns and factor expressions that were largely in accordance with the hypothesized variants. Taken together, our results indicate that variants of posterior cortical atrophy exist but these constitute phenotypical extremes and most individuals fall along a broad clinical-radiological spectrum, indicating that classification into four mutually exclusive variants is unlikely to be clinically useful.

## Introduction

Posterior cortical atrophy is a clinical-radiological syndrome defined by progressive loss of higher-order visual functions, and atrophy that markedly affects posterior brain regions such as the parietal and occipital cortices (Benson *et al*., 1988; Whitwell *et al*., 2007; Koedam *et al*., 2011, Lehmann *et al*., 2011*b*; Crutch *et al*., 2012; Alves *et al*., 2013, Ossenkoppele *et al*., 2015*b*, *a*; Firth *et al*., 2019; Marinescu *et al*., 2019). While multiple pathologies may underlie the posterior cortical atrophy syndrome, the most common biological substrate is Alzheimer’s disease, accounting for ∼80% of the cases (Renner *et al*., 2004; Tang-Wai *et al*., 2004; Montembeault *et al*., 2018). Posterior cortical atrophy has been coined the “visual variant of Alzheimer’s disease” (Kaeser *et al*., 2015) and is the most common non-amnestic variant of Alzheimer’s disease, afflicting ∼5% of all patients diagnosed in a memory clinic setting (Snowden *et al*., 2007). Posterior cortical atrophy manifests at a relatively young age compared to amnestic-predominant Alzheimer’s disease (Mendez *et al*., 2002; Schott *et al*., 2016). Clinically, posterior cortical atrophy can present with a variety of visuoperceptual and visuospatial symptoms, but impairments in other (non-visual) parietal functions (i.e., numeracy and literacy) or primary visual processing (e.g., motion coherence, hue discrimination) are also frequently observed (Crutch *et al*., 2017). Especially in early disease stages, memory, executive functions and language are relatively spared (Schott *et al*., 2016; Crutch *et al*., 2017).

While deficits in visual processing and posterior atrophy are, by definition, the dominant features of posterior cortical atrophy, considerable heterogeneity exists within the clinical and radiological spectrum. This has motivated efforts to categorize this heterogeneity into clinical variants (Crutch *et al*., 2017). The two best characterized variants of posterior cortical atrophy are reflective of the functional organization of the visual system (i.e., ventral and dorsal streams), also referred to as the “what” and “where” pathways (Ungerleider and Haxby, 1994). These variants are called the occipitotemporal (ventral) variant and temporoparietal (dorsal) variant, and are characterized by the presence of prominent visuoperceptual and visuospatial deficits, respectively (Ross *et al*., 1996; Galton *et al*., 2000; Dubois *et al*., 2007; Tsai *et al*., 2011; Huberle *et al*., 2012; Borruat, 2013; Grossi *et al*., 2014). The most recent consensus criteria (Crutch *et al*., 2017) describe two additional variants - a primary visual (caudal) variant, characterized by primary visual processing deficits (Levine *et al*., 1993; Galton *et al*., 2000; Chan *et al*., 2001) and a dominant parietal variant, which presents with prominent non-visual parietal function deficits like dyscalculia, dyslexia and apraxia (De Renzi, 1986; Green *et al*., 1995; Aharon-Peretz *et al*., 1999; Crutch *et al*., 2017). Importantly, characteristics of these posterior cortical atrophy variants are mainly based on single-case studies or studies of limited sample sizes, and several previous attempts to identify consistent clinical and neuroimaging correlates to these variants have failed (McMonagle *et al*., 2006, Lehmann *et al*., 2011*a*; Migliaccio *et al*., 2012). Consequently, in the consensus criteria it is emphasized that current literature provides insufficient cognitive or neuroimaging evidence to support the existence of discrete PCA subtypes and that more research is needed (Crutch *et al*., 2017).

With this in mind, we employed a data-driven Bayesian modelling approach to detect endophenotypes on MRI, and assessed the association between these phenotypes and cognition, in a relatively large sample of well-phenotyped posterior cortical atrophy participants (N=119). The model produces probabilistic atrophy maps for multiple factors (in a continuous fashion) and allows each individual to express each of these atrophy factors to a certain degree. Individual atrophy patterns may encompass multiple brain regions rather than just one single area. This is a more biologically plausible approach than assessing predefined regions-of-interest based on *a priori* classification into mutually exclusive subgroups (i.e., binary/categorical). To test whether the atrophy factor expressions map onto specific visual impairments as proposed in the consensus criteria, we correlated the identified atrophy factors with the four clinical domains of object perception, space perception, non-visual parietal functions and primary visual processing. With this data-driven approach we endeavour to assess how well the reported phenotypical variants capture the clinical and radiological spectrum of posterior cortical atrophy.

## Materials and methods

### Subjects

We selected participants with posterior cortical atrophy from two independent expert centers, the Amsterdam Dementia Cohort of the Amsterdam University medical center (Amsterdam UMC) in Amsterdam, the Netherlands (van der Flier and Scheltens, 2018) and the University of California San Francisco (UCSF) Alzheimer’s Disease Reasearch Center in San Fransisco, CA, USA. All participants underwent dementia screening between June 2000 and July 2017, and inclusion into the present study was based on the following criteria: i) a syndrome diagnosis of posterior cortical atrophy as defined by published diagnostic criteria (Mendez *et al*., 2002; Tang-Wai *et al*., 2004; Crutch *et al*., 2017) and established by consensus in a multidisciplinary meeting, and ii) availability of an MRI scan including a structural volumetric T1-weighted sequence. We excluded participants who had negative biomarkers for amyloid-β pathology [either CSF molecular profile (Zwan *et al*., 2014; Tijms *et al*., 2018) and/or amyloid-PET visual rating (Rabinovici *et al*., 2010; Ossenkoppele *et al*., 2013)]. These criteria resulted in 69 participants from Amsterdam UMC and 50 participants from the UCSF sites. Of these 119 participants, 91 (76%) were amyloid-β positive (40 [34%] on CSF, 28 [24%] on PET and 23 [19%] on both PET and CSF) while for 28 (24%) the amyloid-β status was unknown. We additionally selected 121 amyloid-β negative cognitively normal individuals (age 57.4±8.9, 41%male), who served as a reference group for voxelwise contrasts and were used to contextualize gray matter densities (see “Imaging analyses”). Initial image analyses were performed separately in the two cohorts, but yielded highly similar results. Therefore, the two samples were combined when assessing associations between atrophy and cognition, in order to capitalize on the increased statistical power provided by the larger sample size. The results of the combined cohort are presented in the main text, while the results obtained in the separate cohorts are provided in the supplement. Informed consent was obtained from all participants, and the local medical ethics review committees of the Amsterdam UMC and UCSF approved the study.

### Cognition

Neuropsychological test scores covered two higher-order visual processing domains in both the Amsterdam UMC and UCSF cohorts: Object perception (Amsterdam UMC and UCSF: fragmented letters) and space perception (Amsterdam UMC and UCSF: number location and dot counting) (Boyd *et al*., 2014). The visual test battery administered in the UCSF sample included more tests than in the Amsterdam UMC sample; two additional domains could be assessed within the UCSF cohort only: non-visual dominant parietal functions (calculations, spelling, and reading) and primary visual processing (point location, figure discrimination, shape discrimination, hue discrimination, visual acuity, size discrimination, letter cancellation, static circle detection and motion coherence). Additional neuropsychological test scores covered the following non-visual cognitive domains: memory (Amsterdam UMC: Rey auditory verbal learning test-immediate and delayed recall [15 items/5 trials, Dutch version]; UCSF: California Verbal Learning Test-immediate and delayed recall [9 items/4 trials]), executive functions (Amsterdam UMC and UCSF: Digit-span forwards and backwards; Letter fluency [D]) and language (Amsterdam UMC and UCSF: Verbal fluency [animal naming]) (Kramer *et al*., 2003; Groot *et al*., 2018). Mini-Mental State Examination (MMSE) scores were used as a measure of global cognition.

Before combining neuropsychological data from the two cohorts, all test scores were converted into z-scores using the mean and standard deviation of each separate cohort (to adjust for center effects), and then combined. Furthermore, educational attainment levels were measured using a qualitative scale in the Amsterdam UMC cohort and these were converted to years of education before combining the samples. Cognitive data obtained closest to the date of the MRI scan (up to maximum ± 6 months) were used for the analyses. Availability of cognitive data across neuropsychological tests is presented in table 1.

**Table 1.** Demographic and clinical characteristics. Values depicted are mean±SD, unless otherwise indicated. Differences between groups were assessed using independent samples *t*-tests or Fishers exact-tests, where appropriate. Differences in education were not assessable as education is measured on a qualitative scale at Amsterdam UMC and in years of education at UCSF. Memory test scores (percentage correct) were higher in UCSF but also not directly comparable between the samples as UCSF uses a 9-item test while Amsterdam UMC uses a 15-item test. APOE – Apolipoprotein E, MMSE - mini-mental state examination, RAVLT – Rey auditory verbal learning test, used at Amsterdam UMC, CAVLT – California verbal learning test, used at UCSF a – transformed from a score of 5 on the categorical Verhage scale (Verhage, 1965) b – total words recalled divided by the maximum score possible * - UCSF>Amsterdam UMC # - UCSF<Amsterdam UMC

|  |  | Amsterdam UMC |  | UCSF |  | Combined |  |
| --- | --- | --- | --- | --- | --- | --- | --- |
| N |  | 69 |  | 50 |  | 119 |  |
| Age |  | 62.9±6.1 |  | 66.3±7.7* |  | 63.8±7.1 |  |
| Sex (%male) |  | 41 |  | 34 |  | 38 |  |
| Education, years<sup>a</sup> |  | 11±3 |  | 15±3 |  | 13±4 |  |
| MMSE |  | 20.2±4.7 |  | 20.7±6.2 |  | 20.4±5.4 |  |
| *APOE*ε4, % carriers |  | 55 |  | 41 |  | 50 |  |
| Amyloid-β status,<br>%positive/%unknown |  | 81/19 |  | 70/30 |  | 76/24 |  |
| **Domain** | **Neuropsychological test** | **N** |  | **N** |  | **N** |  |
| Object perception | Fragmented letters (/20) | 37 | 10.3±6.7 | 22 | 9.5±6.5 | 59 | 10.0±6.6 |
| Space perception | Number location (/10) | 39 | 6.6±2.4 | 18 | 3.7±3.5<sup>#</sup> | 56 | 5.6±3.1 |
|  | Dot counting (/10) | 46 | 6.5±2.9 | 21 | 5.5±2.8 | 67 | 6.3±2.9 |
| Non-visual/ dominant<br>parietal | Calculations (/9) |  |  | 13 | 1.6±3.0 |  |  |
|  | Spelling (/20) |  |  | 16 | 12.1±5.4 |  |  |
|  | Reading (/16) |  |  | 14 | 14.6±4.0 |  |  |
| Primary visual<br>processing | Point location (/9.99) |  |  | 9 | 3.0±2.0 |  |  |
|  | Figure discrimination<br>(/20) |  |  | 22 | 16.6±2.8 |  |  |
|  | Shape discrimination<br>(/20) |  |  | 15 | 16.3±3.5 |  |  |
|  | Hue discrimination (/4) |  |  | 22 | 3.1±1.2 |  |  |
|  | Visual acuity (/6) |  |  | 22 | 5.5±1.1 |  |  |
|  | Size discrimination (/1) |  |  | 14 | 0.4±0.4 |  |  |
|  | Letter cancellation (time, s) |  |  | 23 | 96.1±57.0 |  |  |
|  | Static circle detection (/20) |  |  | 15 | 18.7±2.9 |  |  |
|  | Motion coherence (/20) |  |  | 17 | 17.8±3.6 |  |  |
| Memory | RAVLT / CVLT immediate (%correct<sup>b</sup>) | 57 | 31.4±13.2 | 44 | 45.4±18.8* | 101 | 37.5±13.4 |
|  | RAVLT / CVLT delayed (%correct<sup>b</sup>) | 58 | 19.5±20.7 | 44 | 30.1±29.8* | 102 | 24.1±25.5 |
| Executive | Verbal fluency, letter D (correct in 60sec) | 50 | 10.3±4.1 | 43 | 10.1±5.0 | 93 | 10.2±4.5 |
|  | Digit span forward, span (/8) | 56 | 5.2±1.0 | 32 | 5.3±1.2 | 88 | 5.3±1.1 |
|  | Digit span backward, span (/8) | 55 | 3.3±1.0 | 45 | 3.0±1.2 | 100 | 3.1±1.1 |
| Language | Verbal fluency, animal naming (correct in 60sec) | 51 | 13.0±5.3 | 44 | 10.1±5.4<sup>#</sup> | 95 | 11.6±5.5 |

### MRI acquisition

The MR images from Amsterdam UMC were acquired on eight different MRI scanners using previously described standardized acquisition protocols (ten Kate *et al*., 2017) and with a scanner field strength of 1.5T or 3T. The MR images from UCSF were acquired on a 1.5T Magnetom Avanto, a 3T Siemens Tim Trio or a 3T Siemens Prisma Fit scanner. Proportion of participants scanned on a 1.5T scanner were balanced between the two samples, 22% in Amsterdam UMC and 26% in UCSF, and scanner field-strength was used as a covariate in all imaging analyses.

### Imaging analyses

Image processing steps were performed separately for the Amsterdam UMC cohort, the UCSF cohort and the combined cohort. T1-weighted images were segmented into gray matter, white matter and CSF volumes using statistical parametric mapping version 12 (Wellcome Trust Centre for Neuroimaging, UCL, London, UK). Diffeomorphic anatomical registration through exponentiated Lie algebra (Ashburner, 2007) was then used to generate a study-specific template by aligning gray matter images nonlinearly to a common space. Gray matter images were spatially normalized to the study-specific template using individual flow fields. Modulation was applied to preserve the total amount of signal, and images were smoothed using an 8mm full-width-at-half-maximum isotropic Gaussian kernel. The resulting gray matter density images were used to assess the whole-brain spatial distribution of atrophy by performing voxelwise contrasts between participants with posterior cortical atrophy and controls. Next, the gray matter density images were converted into W-score maps (i.e., control-normalized z-scores adjusted for covariates) (La Joie *et al*., 2012, Ossenkoppele *et al*., 2015*a*; van Loenhoud *et al*., 2017) by performing voxelwise standardization to the control group, regressing out the effects of age, sex, intracranial volume, scanner field strength and whole-brain gray matter atrophy (operationalized as gray matter to intracranial volume ratios). The resulting W-maps were log-transformed and used as the input to the Bayesian modelling framework (Fig. 1).

**Figure 1.**
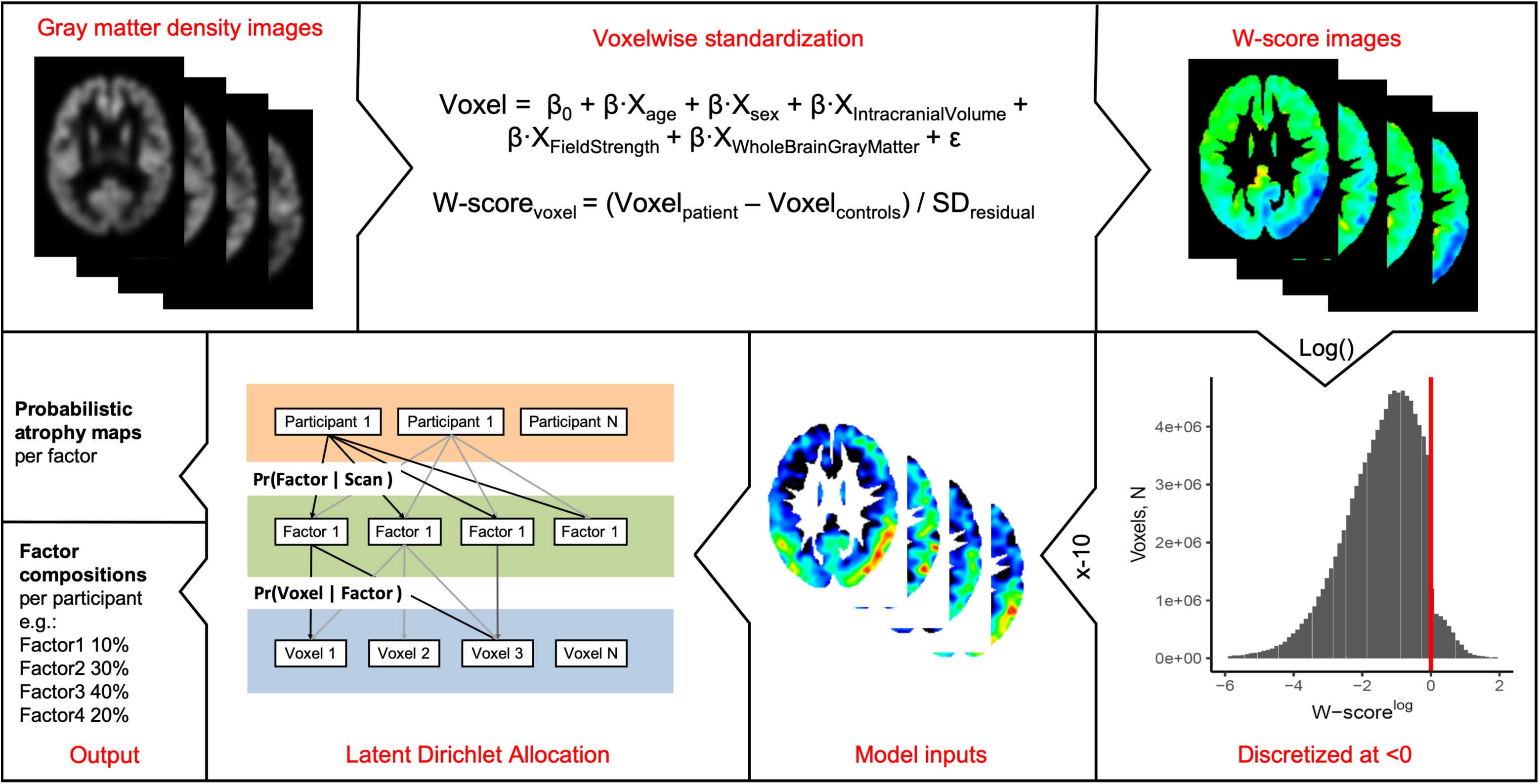
A Bayesian model of participants, atrophy factors, and structural MRI. Each participant expresses one or more factors to a certain degree and each factor is associated with distinct but possibly overlapping patterns of brain atrophy. The estimated parameters are the probability that a participant (scan) expresses a particular factor [i.e., Pr(Factor | Scan)] and the probability that a factor is associated with atrophy at an MRI voxel [i.e., Pr(Voxel | Factor)]. Adjusted with permission from Zhang et al 2016.

### Bayesian modelling

We employed a Bayesian modelling approach called latent Dirichlet allocation (LDA) to discover atrophy patterns that covary across participants in order to identify latent atrophy factors present within our sample of posterior cortical atrophy participants. This method has been adapted for structural MRI data in a previous study including patients with Alzheimer’s disease (Zhang *et al*., 2016). The LDA model considers each scan as an unordered collection of voxels associated with a predefined number of latent atrophy factors (*K*). The LDA model assumes that a scan is summarized by the degree of atrophy (derived from the W-scores described above) in each voxel of the scan, and allows each individual’s scan to be associated with multiple factors and each factor to be associated with multiple voxels. More specifically, given a dataset of scans, the algorithm estimates the probability of atrophy at a particular voxel given a factor [Pr(Voxel | Factor)] and the probability that a factor is associated with a particular scan [Pr(Factor | Scan)] (Fig. 1). To achieve these estimations, the continuous log-transformed W-score images were first discretized so that a W-score of <0 at a given voxel of a particular scan would imply above-average atrophy at the voxel relative to the controls, adjusted for the effects of age, sex, intracranial volume, scanner field strength and whole-brain atrophy. W*-*scores >0 were set to zero (values above 0 reflect gray matter density greater than the control group). Then, the W-scores were multiplied by −10 and rounded to the nearest integer, so that larger positive values indicated more severe atrophy. Given the discretized voxelwise atrophy of the posterior cortical atrophy participants and the number of latent atrophy factors *K*, the variational expectation maximization algorithm (www.cs.princeton.edu/~blei/lda-c/) was applied to estimate Pr(Factor | Scan) and Pr(Voxel | Factor). A latent factor (Pr[Voxel | Factor]) can be visualized as a probabilistic atrophy map. Pr(Factor | Scan) is a probability distribution over latent atrophy factors, representing the factor composition of the participant (scan). For example, when four factor expressions are estimated for a model with four factors (*K*=4), Pr(Factor | Scan) might be: 10% factor 1, 30% factor 2, 40% factor 3 and 20% factor 4. These factor compositions add up to 100%, and the individual components will henceforth be referred to as (atrophy) factor expressions, while the combination of the factor expressions constitutes an individual’s factor composition. Because the factor expressions add up to 100%, an individual’s expression of a particular factor could be regarded as the proportion of atrophy falling into a specific (but not necessarily localized) anatomical region rather than in the anatomical regions encompassed by the other factors. Therefore, factor expressions and factor compositions are reflective of an individual’s spatial distribution of atrophy rather than its severity.

The algorithm was rerun with 20 different random initializations, and the solution with the best model fit (based on log-likelihood) was selected. The random initializations led to highly similar solutions. Sixty iterations were run for each random initialization, although each run plateaued after around 30-50 iterations (Supplemental Fig. 1). An important model parameter is the number of latent factors (*K)*. We ran models allowing for 2 to 6 factors (*K=*2-6) but we will focus on the results obtained by the model that allows for four factors (*K*=4) in the main text, in accordance with the number of posterior cortical atrophy variants proposed in the diagnostic criteria (i.e., dorsal, ventral, caudal and dominant parietal variant) (Crutch *et al*., 2017). Results from the other models (i.e. *K* = 2, 3, 5 and 6) will be presented in the supplement.

### Statistical analyses

Statistical analyses were performed using statistical parametric mapping version 12 and R version 3.5.2. To assess the cross-sectional associations between atrophy factor expressions and cognition, we used multiple linear regression analyses, adjusted for education, using the “lme4” package in R. Note that the factor expressions were already adjusted for age, sex, whole-brain atrophy, intracranial volume and scanner field-strength effects in the LDA model (see “Imaging analyses” and “Bayesian modelling” sections). In the *K*=4 models outlined in the main text, we included three of the four factors in the predictor set and the fourth was implicitly modelled because factor expressions of the four factors add up to 100%. The model was therefore: y=β_0_ + β_1_·*K*_1_ + β_2_·*K*_2_ + β_3_·*K*_3_ + β_education_·education + ε, with y denoting cognition, β the regression coefficients, *K* the atrophy factor expressions and ε the residual. The relative effects of the three directly modelled factors were calculated using the implicitly modelled fourth factor (β_0_) as a reference. All models were repeated using a different atrophy factor implicitly modelled to obtain pairwise differences for all factor comparisons. The same approach was used to assess the factor expressions obtained by the *K* = 2, 3, 5 and 6 models, results of which are provided in the supplement (Supplemental Fig. 2).

The association between a single factor expression and cognition obtained by the regression models should be interpreted based on the characteristics of the factors obtained by the LDA model. Factor compositions represent the proportion of atrophy falling into a specific region rather in the others. By using one factor as the reference in each model, effects obtained represent the association of the factor compared to the reference factor in each pairwise comparison. A negative association of factor X with cognition Z would therefore state that individuals with a greater proportion of atrophy in regions associated with factor X, rather than factor Y, have worse scores on domain Z. All results regarding the factor *vs* cognition associations represent these pair-wise comparisons (i.e. K1-K2, K1-K3, K1-K4, K2-K3, K2-K4, K3-K4). Statistical significance for all models was set at α=0.05 and we performed *post-hoc* adjustment for multiple comparisons using the false-discovery-rate (FDR) method, correcting for a total of 132 comparisons based on the number of pairwise comparisons (6) times the number of tests (22). Both uncorrected and FDR-corrected results are presented.

### Data availability

The code for the Bayesian modelling approach is publicly available at (https://github.com/ThomasYeoLab/CBIG/tree/master/stable_projects/disorder_subtypes/Zhang2016_ADFactors). Data used in the present study may be available upon request to the corresponding author.

## RESULTS

Demographic and clinical characteristics of the Amsterdam UMC, UCSF and combined samples are presented in Table 1. Mean age of the total sample was 63.8±7.1, 38% were male and MMSE was 20.5±5.2. Voxelwise contrasts compared to controls revealed a classical posterior cortical atrophy pattern for both the Amsterdam UMC and UCSF cohorts, covering the middle and inferior temporal gyrus, inferior and medial parietal areas and the occipital cortex (Fig. 2). The atrophy pattern was slightly lateralized to the right-hemisphere, and very similar across the two cohorts. This similarity between cohorts was also reflected in the voxelwise contrasts of the combined sample compared to controls.

**Figure 2.**
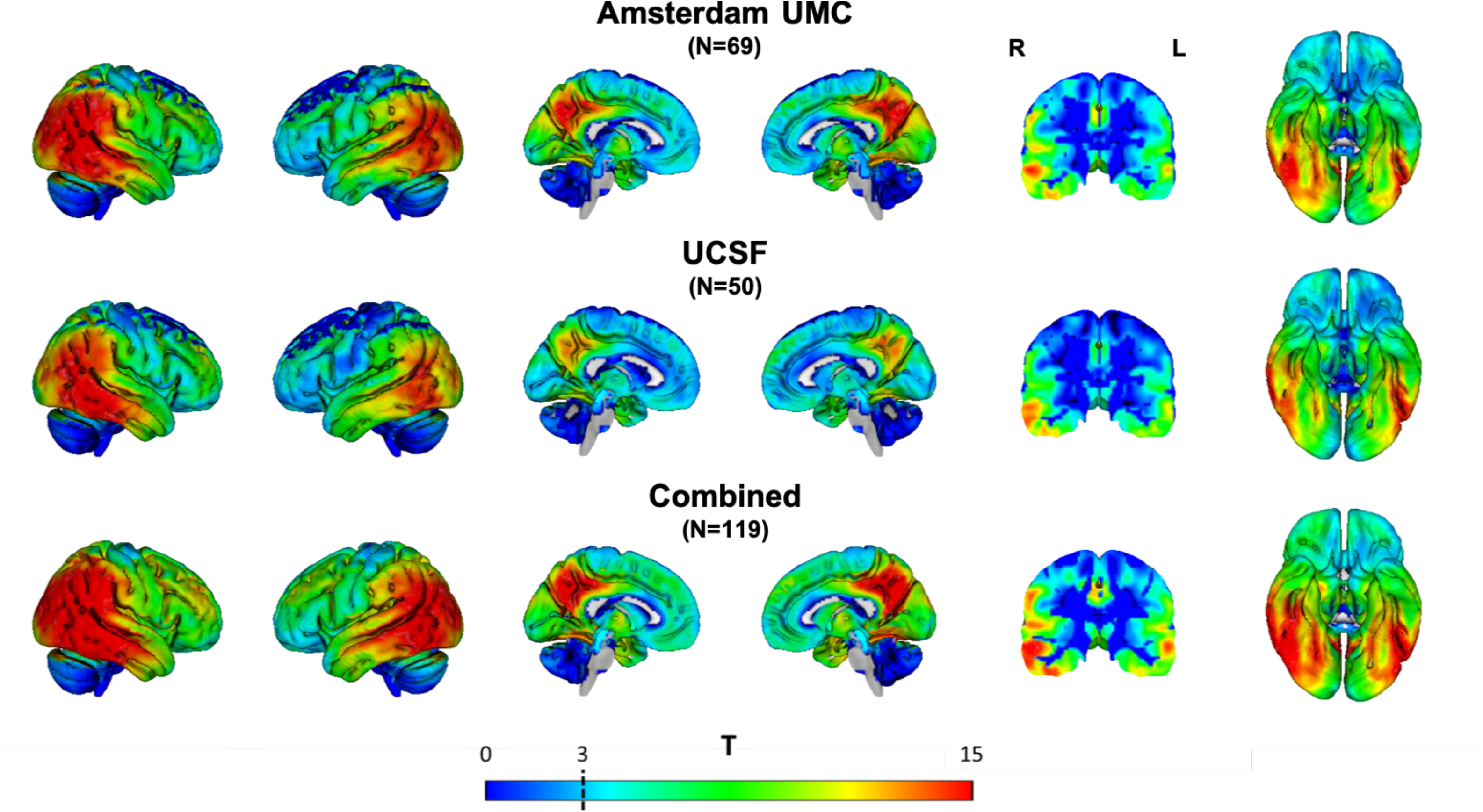
Exploratory voxel-wise contrasts between posterior cortical atrophy participants and controls. Voxelwise T-maps are adjusted for the effects of age, sex, intracranial volume, whole-brain atrophy and scanner field strength. Significant voxels at T>3.

### Latent atrophy factors

The Bayesian model (*K* = 4) revealed four distinct but partially overlapping latent atrophy factors (Fig. 3), which were similar in the Amsterdam UMC and UCSF cohorts. The first factor (“right-dorsal”) included the right lateral temporoparietal cortex as well as bilateral medial parietal regions. The second factor (“right-ventral”) included the right medial and lateral occipital cortex, extended inferiorly into the temporal cortex, and also covered part of the inferior parietal cortex. The third factor (“left-ventral”) included the left medial and lateral occipital cortex, inferior temporal cortex, and inferior parietal cortex. The fourth factor (“limbic”) mainly included bilateral medial-temporal areas as well as medial frontal regions (Fig. 3). Since factors were, overall, very similar between the two cohorts and VBM analyses also produced similar atrophy patterns (Fig. 2), the results of the combined factor are highlighted in the rest of the main text, while results from the two separate cohorts are provided in the supplement (Supplemental Fig. 3 and 6).

**Figure 3.**
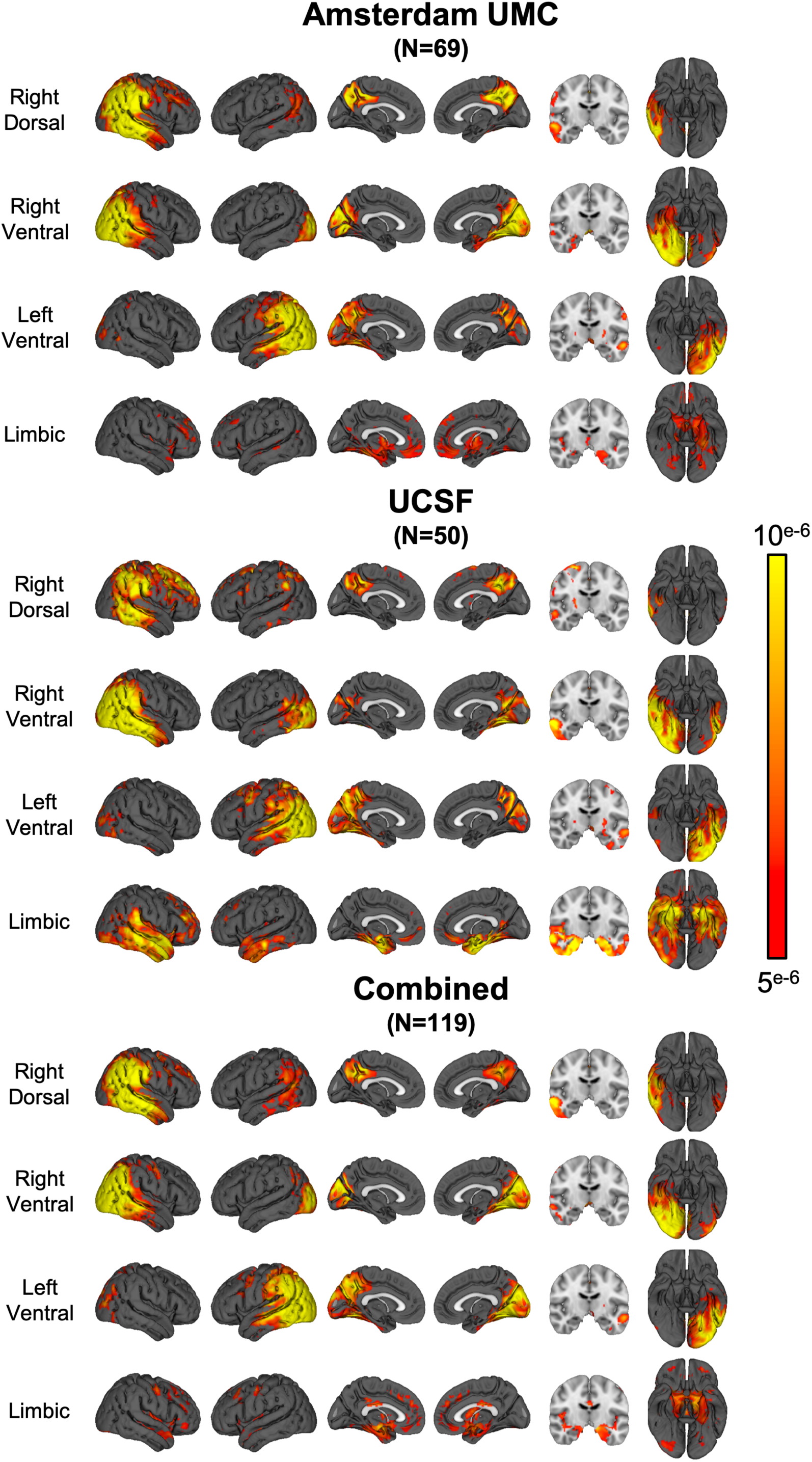
Atrophy factors revealed by latent Dirichlet allocation (*K*=4) Intensity of voxels signify the probability [Pr(Voxel | Factor)] of a voxel belonging to one of the four factors. Scale is truncated at [Pr(Voxel | Factor)]=5^e-6^ for visualization purposes.

### Individual factor compositions

Factor compositions of the combined sample reveal that the majority of posterior cortical atrophy participants expressed a combination of multiple atrophy factors rather than predominantly expressing only one of the factors (Fig. 4). This indicates that most participants have atrophy that extends across multiple regions rather than focal atrophy confined to a single region. A similar distribution was observed when we stratified individuals according to clinical disease severity (MMSE: 30-24 *vs* 23-18 *vs* 17-6; Supplemental Fig. 4). To assess whether factor expressions were partly driven by global atrophy, we examined the relationship between factor expressions and whole brain gray matter to intracranial volume ratios. We observed a significant correlation only between the limbic factor and whole-brain gray matter to intracranial volumes ratios (lower values indicate more atrophy; r=-0.43, p<0.001), while the other factors did not show a correlation (range: r=0.11 to 0.19, all p>0.05). This indicates that individuals with a higher proportion of atrophy in the limbic factor when compared to the other factors, tend to have more atrophy overall. Furthermore, we observed a significant correlation only between the right-dorsal factor and age (r=-0.26, p=0.005), which indicates that individuals with a higher proportion of atrophy occurring in the right-dorsal factor (compared to the others) tend to be younger. There were no associations between factor expressions and sex (range: t=-1.51 to 1.75, all p>0.05), APOEε4 (+/-; range: t=-1.28 to 1.00, all p>0.05) or handedness (right/non-right handed; range: t=-0.36 to 1.31, all p>0.05).

**Figure 4.**
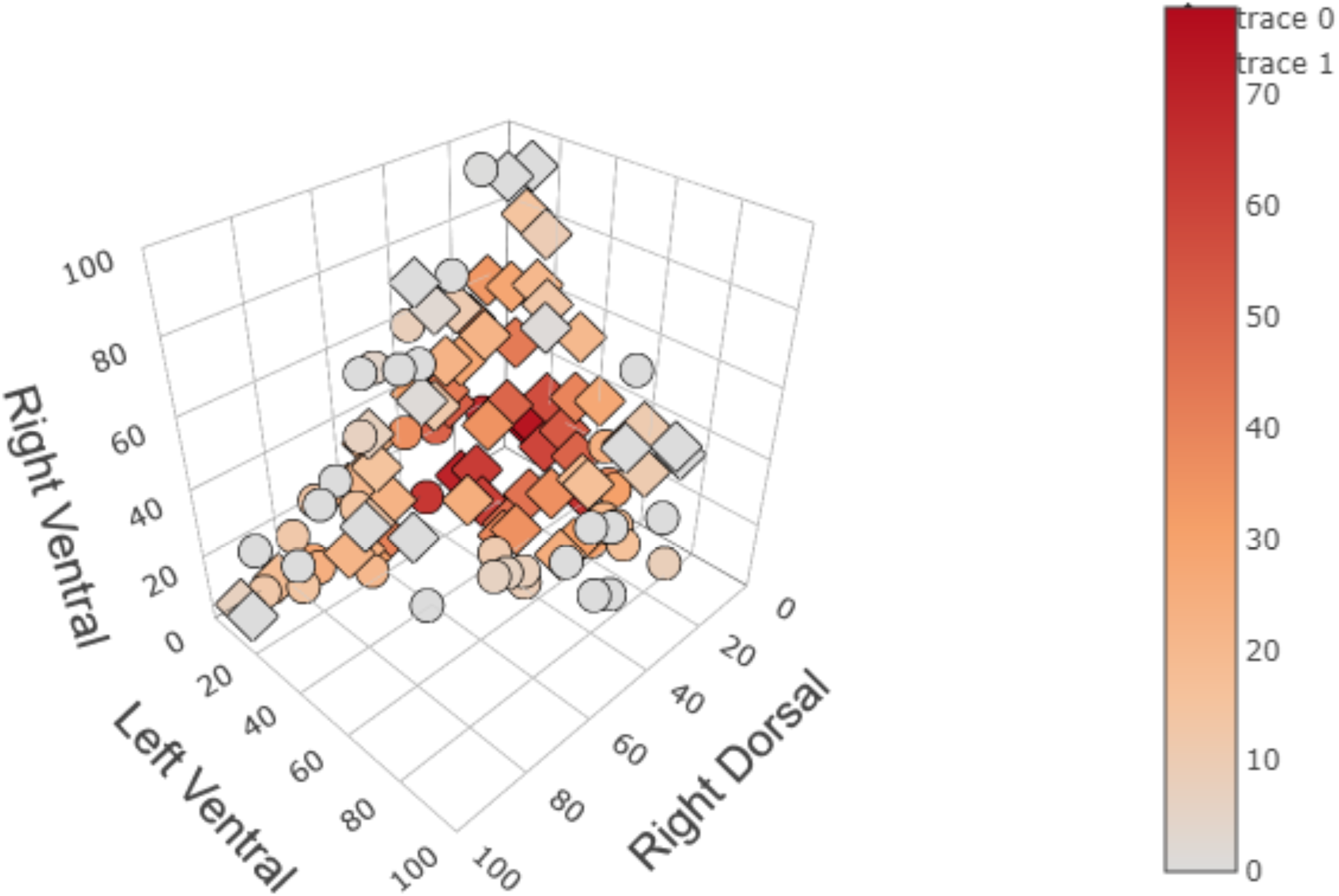
Atrophy factors compositions for the combined sample. This 4D plot displays the factors right-dorsal, left-ventral and right-ventral on the x, y and z axes and the limbic factor is displayed by the color gradient of the markers. Displayed factor compositions are for the combined sample and each marker represents one participant, the Amsterdam UMC participants are denoted by the diamond shaped markers while the UCSF participants are displayed with circles. Expressions of the four factors adds up to 100%.

### Associations between factor expression and higher-order visual processing

We assessed the associations between factor expression and higher-order visual processing (i.e., object and space perception), which are highly relevant to the clinical presentation of posterior cortical atrophy. Object perception was assessed by the fragmented letter test and our results indicate that object perception is associated with atrophy that predominantly affects the right-ventral and limbic regions. Specifically, we observed that right-ventral factor expression was negatively associated with fragmented letter scores compared to right-dorsal and left-ventral factor expressions (β=-0.35, p=0.008 uncorrected; β=-0.48, p=0.001 FDR-corrected). Limbic factor expression was also associated with worse fragmented letters scores compared to left-ventral factor expression (β=-0.34, p=0.043 uncorrected; Fig. 5 and Supplemental Table 1).

**Figure 5.**
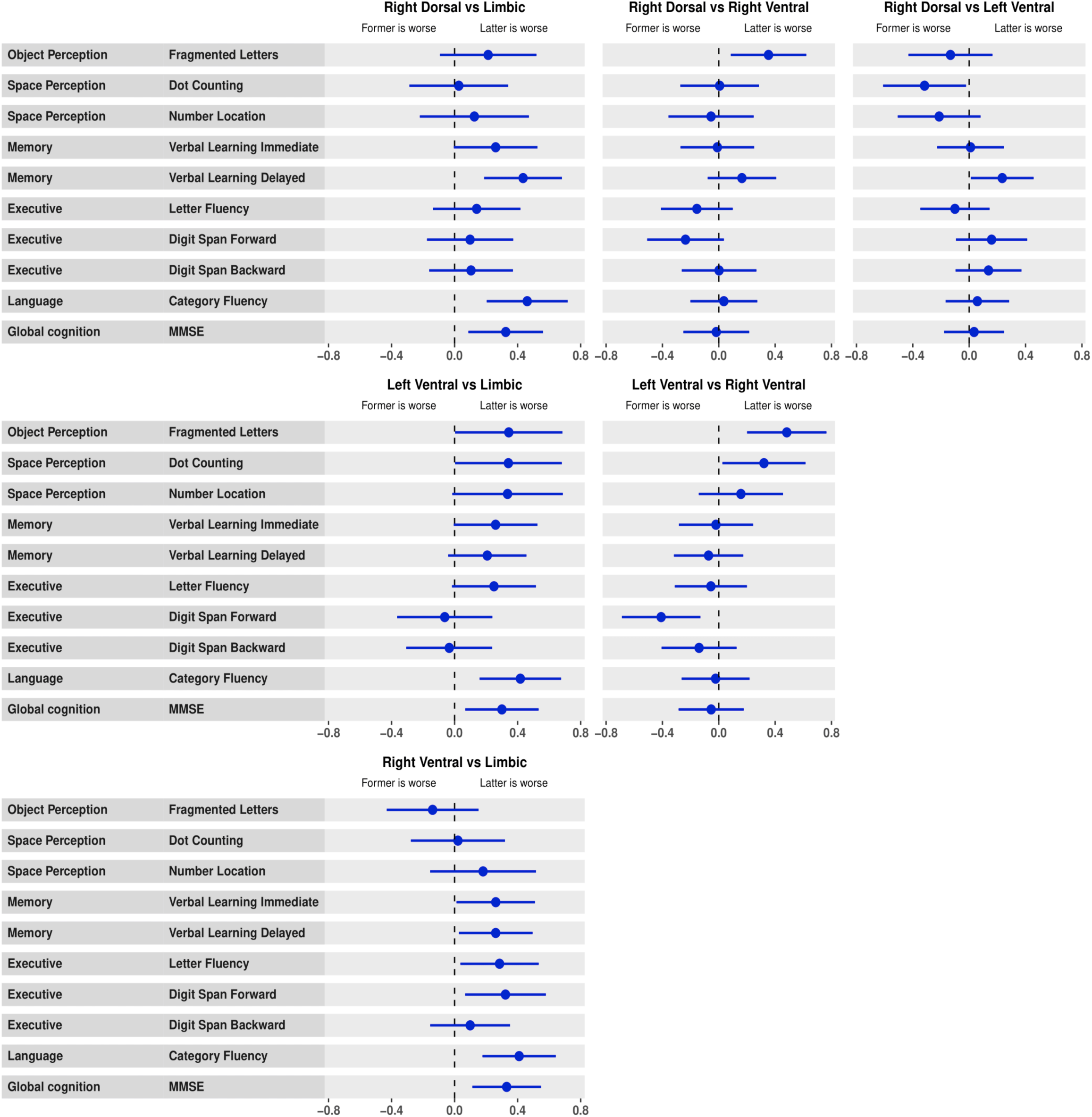
Associations between factor expressions and neuropsychological tests in the combined sample. The plot contains relative cross-sectional effects from linear regression models. Lines indicate the 95% confidence intervals and a significant effect (uncorrected for multiple comparisons) is denoted by confidence intervals not including x=0.

Space perception was assessed using the dot counting and number location tests and our results indicate that space perception is associated with atrophy that predominantly affects right-dorsal, right-ventral and limbic regions (as opposed to left-ventral). Specifically, right-ventral, limbic and right-dorsal factor expression were negatively associated with dot-counting compared to the left-ventral factor (β=-0.32, p=0.030 uncorrected, β=-0.34, p=0.044 uncorrected, β=-0.32, p=0.031 uncorrected). This same pattern was observed for number location scores, although none of the effects reached statistical significance (Fig. 5 and Supplemental table 1; Fig. 5 and Supplemental Table 1).

### Associations between factor expression and non-visual dominant parietal and primary visual processing functions

Primary visual processing was negatively associated with right-ventral compared to left-ventral (hue discrimination: β=-0.59, p=0.048 uncorrected), limbic (letter cancellation: β=-0.59, p=0.028 uncorrected) and right-dorsal factor expression (shape discrimination: β=-0.59, p=0.027 uncorrected; Supplemental Table 1 and Supplemental Fig. 5). This indicates that individuals with atrophy that predominantly affects the right-ventral factor tend to have worse performance on primary visual processing.

With regard to non-visual parietal functions, we observed a trend towards worse calculations and spelling scores in the right-dorsal, left-ventral and right-ventral factors, compared to limbic. These findings suggest that non-visual “parietal’ functions are associated with extra-limbic factors (Supplemental Table 1 and Supplemental Fig. 5).

### Associations between atrophy and MMSE, memory, executive and language functioning

Beyond the visual processing domains, we examined the association between factor expressions and memory, executive and language functions, as well as global cognition measured by MMSE. Across verbal learning, letter fluency, digit span, and category fluency tests, we found negative association with the limbic factor loading compared to the other factors. For MMSE, we also found more associations with limbic factor expression compared to the other factors, while associations between the extra-limbic factors were sparse (Fig. 5 and Supplemental Table 1). These findings suggest that individuals with a higher proportion of atrophy in the limbic regions, as opposed to the other (neocortical) regions, tend to have worse performance on non-visual cognitive functions, and worse global cognition.

### Case series of participants corresponding to distinct posterior cortical atrophy variants

While associations between factor expressions and cognition revealed relationships that are largely in accordance with established brain-behavior relationships, individual factor compositions indicated that the vast majority of participants express atrophy across multiple factors rather than in one primarily. This suggests that individual atrophy patterns span across multiple brain networks and factors; our results therefore do not support the notion that discrete phenotypical variants of posterior cortical atrophy are common. To provide an explanation for the description of these variants in earlier studies, we include a case description of four participants who were selected based on an isolated relative impairment in one of the cognitive domains most relevant to posterior cortical atrophy: object perception, space perception, non-visual/dominant parietal functions or primary visual processing (Fig. 6A). These scores were obtained by averaging the scores across neuropsychological tests within each domain. From this plot it is evident that, similar to what we observed for the factor compositions, most participants have impairments across multiple cognitive domains, and only a few had a clinical phenotype that was characterized by isolated impairments (see annotated markers in Fig. 6A). We outlined the clinical and radiological characteristics of these four cases in Fig.s 6B and C. Case 1 was a 69-year-old female (MMSE: 25) with pronounced object perception impairment and prominent, right-lateralized, inferior parietal and occipitotemporal atrophy, extending towards the inferior temporal lobe. The clinical and radiological phenotype of this case is compatible with the occipitotemporal (“ventral”) variant of posterior cortical atrophy described in literature, and this participant mainly expressed the right-ventral factor (80% loading). Case 2 was a 60-year-old female (MMSE: 24) with pronounced space perception deficits and atrophy mainly in the right occipital, parietal and temporal cortices but not in the inferior temporal lobe. This case also showed atrophy in right dorsolateral-prefrontal areas. This phenotype matches with the temporoparietal (“dorsal”) variant, and this participant had a relatively high right-dorsal factor expression (50% loading). Case 3 was a 76-year-old male (MMSE: 10) with low scores on non-visual, dominant parietal functions but also on the other three domains.

**Figure 6.**
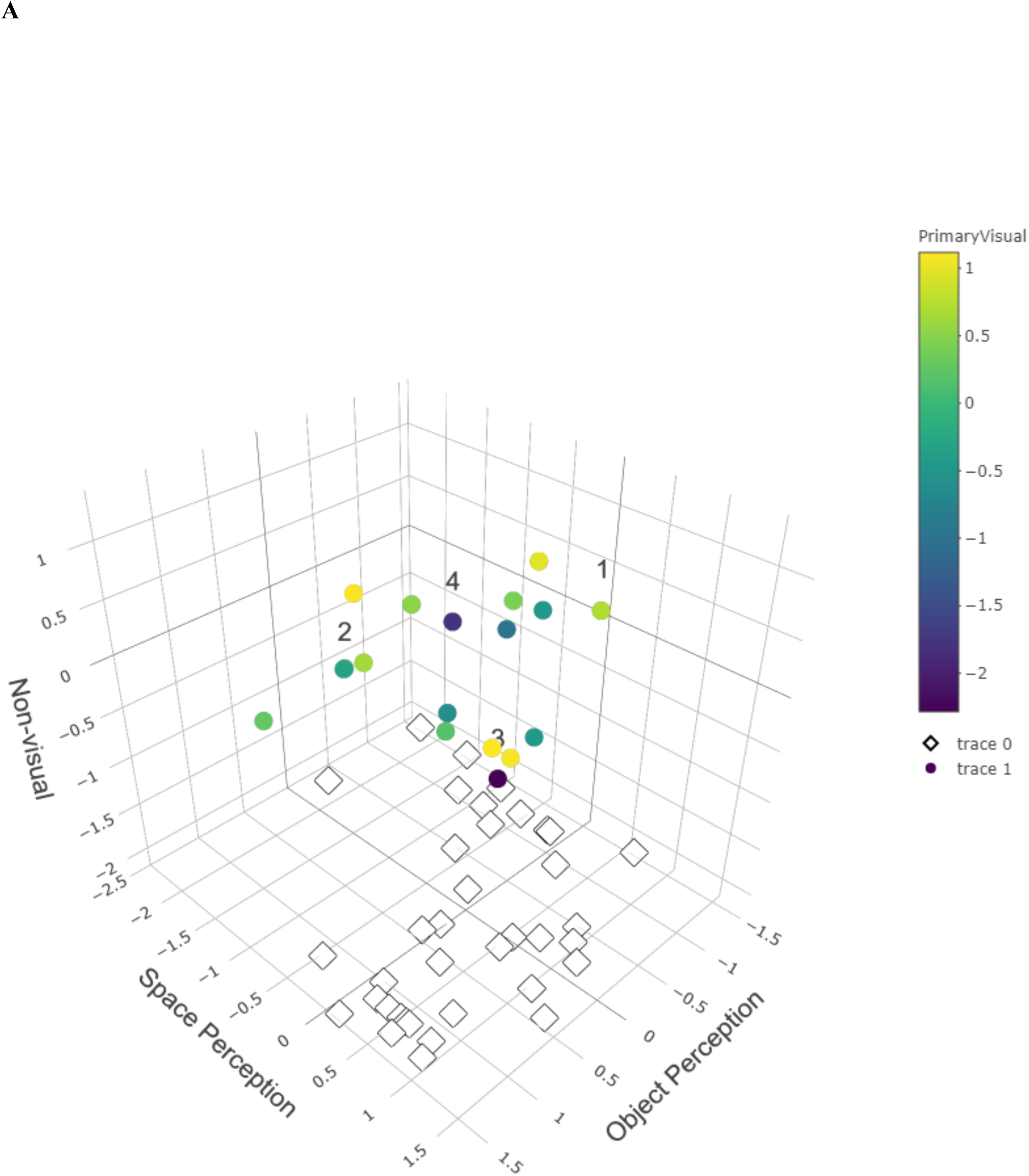
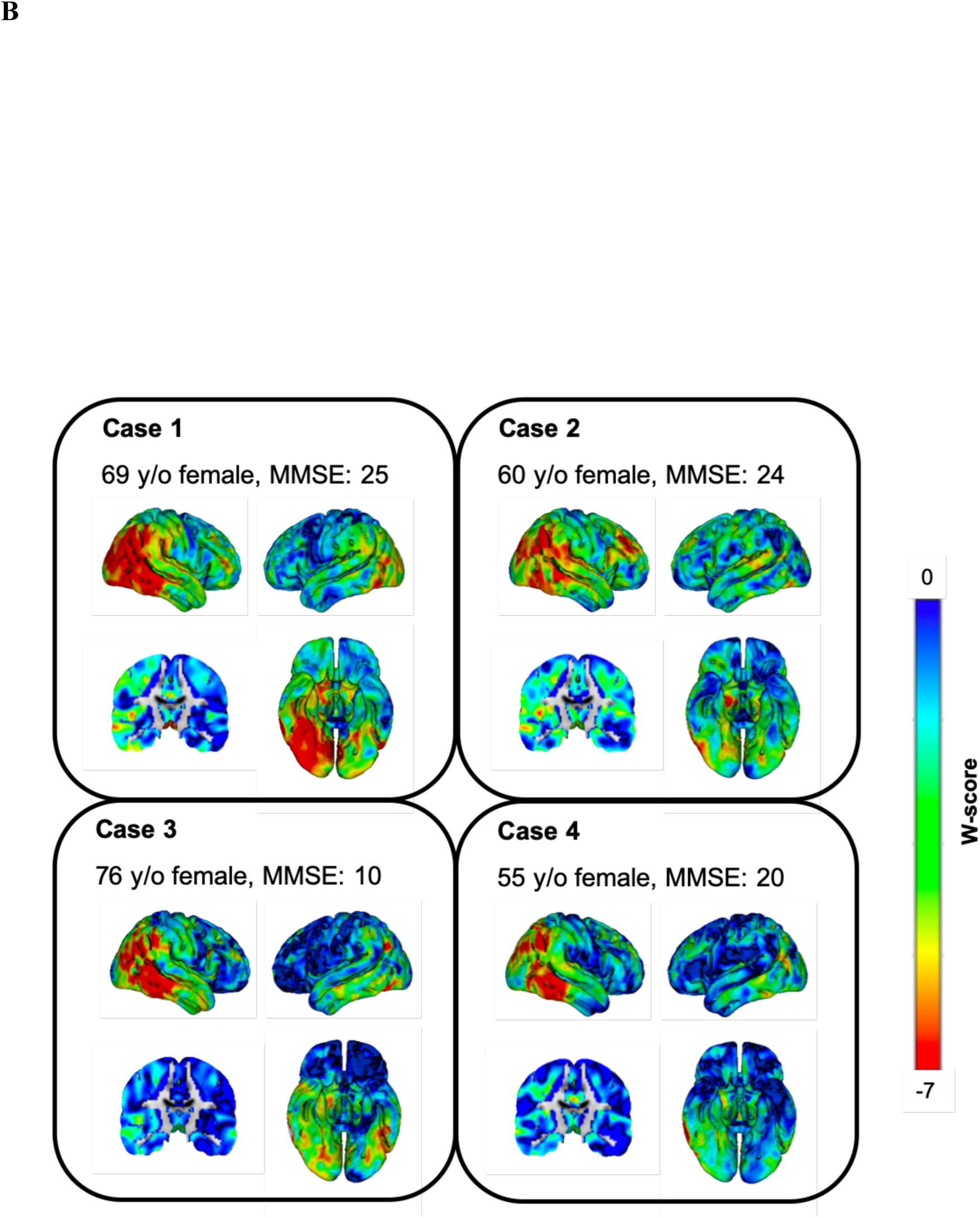
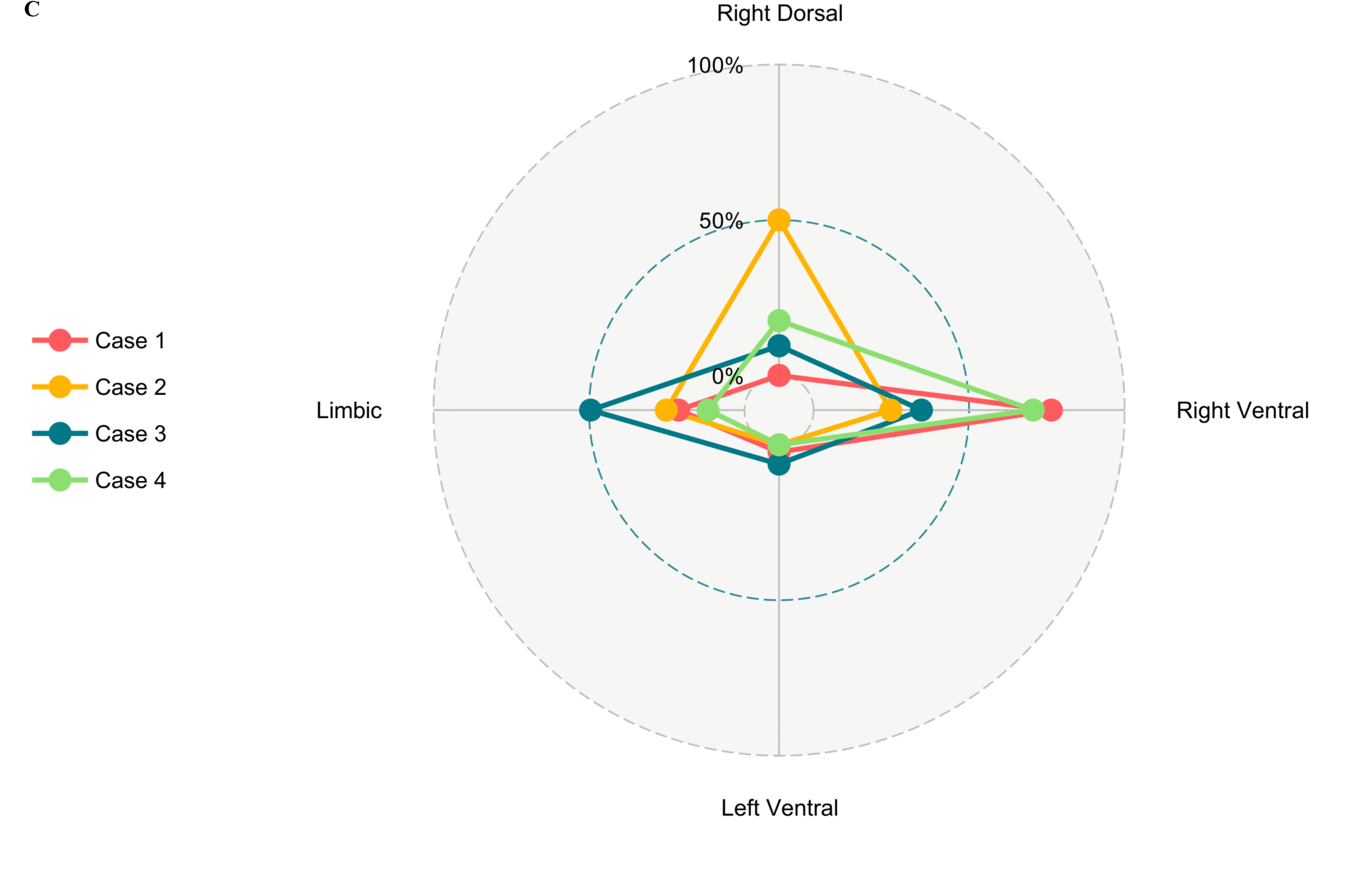
Case series of extreme clinical phenotypes. (A) This 4D plot displays scores on the object perception, space perception and non-visual/dominant parietal domains on the x, y and z-axes. Primary visual processing scores are displayed by the color gradient of the markers. As the Amsterdam UMC cohort did not include any non-visual/dominant parietal or primary visual processing tests, these scores are projected onto the x and y-axes, and colorless. We selected cases with isolated relative impairments, one for each domain. Selected cases within this distribution are annotated by number 1 through 4, only participants with scores on all four domains (from the UCSF sample) were eligible for selection. (B) Displays the clinical characteristics of the four selected cases as well as the regional spread of atrophy indicated by W-scores. Lower W-scores represents more atrophy. (C) This radarplot displays individual factor expressions of the four selected cases.

Radiologically, this case presented with a right-lateralized temporoparietal atrophy pattern and factor expression was mainly in the limbic factor (50% loading). The phenotype of this participant fits the clinical description of the dominant-parietal variant but less clearly the radiological phenotype. Finally, case 4 was a 55-year-old female (MMSE: 20) who had low primary visual processing scores and atrophy that was mainly localized in the right inferior parietal cortex and posterior part of the lateral temporal cortex, with a high right-ventral factor expression (70% loading). This phenotype matches the clinical description of the caudal (or primary visual) variant in literature but the atrophy pattern was not markedly caudal. (Fig. 6B-C). These observations indicate that while the clinical phenotypes are recognizable among a sample of posterior cortical atrophy participants, these are the exceptions rather than the rule, and even than do not uniquely map 1 on 1.

## Discussion

In the present study, we employed a data-driven approach to identify phenotypical variants of posterior cortical atrophy by detecting latent atrophy factors and assessing the associations between these factors and cognitive domains known to be affected in posterior cortical atrophy (i.e., object perception, space perception, non-visual dominant parietal and primary visual processing). We included independent cohorts from two expert centers, which were combined into a large sample of this relatively rare condition. As expected, voxel-based morphometry confirmed a characteristic posterior atrophy pattern in the whole sample, with a slight lateralization to the right hemisphere. A Bayesian modelling framework was used to detect atrophy patterns that covary across participants and identified four distinct but partially overlapping atrophy factors; right-dorsal, right-ventral, left-ventral and limbic. When we evaluated these atrophy factors in our patients, we observed that the vast majority expressed multiple factors rather than primarily expressing only a single factor. With regards to associations between the atrophy factors and higher-order visual processing, we found that object perception was associated with atrophy that predominantly affects the right-ventral and limbic regions. Furthermore, space perception was associated with atrophy that predominantly affects right-dorsal, right-ventral and limbic regions (compared to left-ventral). Primary visual functions were also associated with atrophy that predominantly affects the right-ventral factor but we found no associations for dominant-parietal functions. These findings indicate that atrophy patterns within subjects were associated with particular cognitive functions, mostly in line with known brain-behavior relationships. However, similar to expressions across atrophy factors, scores across cognitive domains revealed that most participants had impairments on multiple visual processing and non-visual parietal functions, rather than being primarily impaired in one. Four participants selected based on a relative impairment on a single domain revealed individual atrophy patterns that were largely in accordance with the hypothesized variants of posterior cortical atrophy, but these cases constituted the exception rather than the norm and even then where not mutually exclusive. Taken together, our Bayesian modelling approach captures atrophy factors that are in accordance with the most well-described phenotypical variants of posterior cortical atrophy (i.e., dorsal and ventral variants) and these brain regions are individually associated with specific clinical features. However, participants display a constellation of affected brain regions and symptoms, and individual phenotypes are intricate. Therefore, classification into four overarching phenotypes is unlikely to be clinically useful.

### Clinical and neurobiological heterogeneity within the spectrum of posterior cortical atrophy

Our results are in line with previous studies with more limited sample sizes that have tried to identify posterior cortical atrophy variants using neuroimaging techniques (Lehmann *et al*., 2011*a*; Migliaccio *et al*., 2012). One study stratified participants into object and space perception subgroups according to relative impairments on these domains and looked at differences in regional cortical thickness (Lehmann *et al*., 2011*a*). This study did find trends towards differences in regional cortical thickness, with the space perception subgroup showing thinner cortex in focal dorsal areas (e.g., superior parietal cortex) and the object perception group in focal ventral areas (e.g., inferior temporal lobe), but these differences were subtle and there was substantial anatomical overlap between the subgroups. This led to the conclusion that there was insufficient indication for the existence of distinct posterior cortical atrophy variants. In the present study, we found that the right-ventral atrophy factor was negatively associated with object perception compared to the right-dorsal factor but dorsal *vs*. ventral associations were not detected with regard to space perception. We did observe that both object and space perception, as well as primary visual functions, were associated with the right-ventral and right-dorsal factors compared to the left-ventral factor. While higher-order visual processing is not clearly lateralized (Ungerleider and Haxby, 1994), it has consistently been found that posterior cortical atrophy presents with a tendency towards right-lateralized atrophy (Migliaccio *et al*., 2009; Crutch *et al*., 2012; Lehmann *et al*., 2013). Since visual processing impairments are the hallmark feature of posterior cortical atrophy, a link to vulnerability of the right-hemisphere is conceivable. Another neuroimaging investigation of posterior cortical atrophy employed diffusion-tensor imaging and assessed tractography measures and cognition. This study found that all investigated participants had ventral white matter tract abnormalities (e.g., inferior longitudinal fasciculus), while only some had additional dorsal stream abnormalities (Migliaccio *et al*., 2012), suggesting that ventral pathways might be affected more than dorsal pathways in posterior cortical atrophy. The neuropsychological assessment of participants in this study also revealed that ventral (object perception) symptoms were present in all participants (Migliaccio *et al*., 2012), which is in contrast to two other studies that observed that dorsal (space perception) symptoms were more prevalent and that ventral (object perception) symptoms only become evident in later stages (McMonagle *et al*., 2006) or present in addition to dorsal symptoms (Tsai *et al*., 2011). In the present study, we found that impairments in object and space perception across the sample were balanced, which is in line with the investigation by previous work wherein the object and space perception subgroups were about equal in size (Lehmann *et al*., 2011*a*).

Deficits in visual processing functions are, by definition, the most prominent features of posterior cortical atrophy, but memory, executive and language functions impairment are also often observed, although these impairments – especially language (McMonagle *et al*., 2006; Crutch *et al*., 2013; Magnin *et al*., 2013; Schott and Crutch, 2019) – are often only present in the later stages of the disease (Schott *et al*., 2016; Crutch *et al*., 2017). We found that individuals for whom the majority of their atrophy was specific to limbic regions had worse memory, executive and language scores compared to those for whom the majority of the atrophy took place in the extra-limbic factors. It has been reported in a previous study that performance on verbal learning is associated with volume of the inferior parietal lobules in posterior cortical atrophy (Ahmed *et al*., 2018), rather than the medial temporal lobe, which contrasts with our findings. However, individuals with atrophy that predominantly affected the limbic regions also showed worse global cognition, indicating that disproportionate limbic atrophy indicates worse cognition overall. As this factor was also the only one associated with global atrophy, it seems that high limbic factor expression might be a feature of late-stage posterior cortical atrophy, which is in accordance with findings reported in previous studies (Lehmann *et al*., 2012; Firth *et al*., 2019; Phillips *et al*., 2019).

### A case against classification of posterior cortical atrophy into distinct variants

Classical neuroscientific literature describes the ventral and dorsal pathways of visual processing, which together make up the model of the “two-streams hypothesis” and are sometimes called the “what” and “where” pathways (Ungerleider and Haxby, 1994). These processing streams respectively encompass occipitotemporal and temporoparietal areas, and one may assume that atrophy in one of these regions lead to specific clinical phenotypes in posterior cortical atrophy. In the present study we found three distinct (although partly overlapping) atrophy factors which roughly corresponded to the ventral and dorsal visual processing pathways, namely the right and left-ventral factors, and the right-dorsal factor. We found that right-ventral factor expression was associated with object perception compared to right-dorsal or left-ventral, while space perception was associated with both right-ventral factor and right-dorsal expression compared to left-ventral. These associations are therefore largely in line with the “two-streams hypothesis”, although space perception was not discretely associated with the (right-)dorsal factor. It might have been that our method was not able to accurately delineate this association, but a previous study implementing the same approach to structural MRI data in an mild-cognitive impairment and Alzheimer’s disease dementia population did find distinct brain-behavior associations in a biologically plausible manner (Zhang *et al*., 2016). Moreover, previous examinations that have focused on dorsal *vs.* ventral neuroimaging features and clinical symptoms (Lehmann *et al*., 2011*a*; Migliaccio *et al*., 2012) have been unable to provide definitive results. The explanation for this may lie in the fact that individuals do not exclusively express atrophy in either the dorsal or the ventral regions, as illustrated by the factor compositions in the present study. Moreover, clinical impairments are also not limited to a single domain but spread across multiple domains. This combination indicates that the dorsal and ventral stream variants are either too rare, or too much overlapping to be discernable.

Evidence for the existence of the other two, admittedly less well-defined, variants of posterior cortical atrophy described in literature (i.e., the caudal and dominant parietal variants) is even more limited. The dominant parietal variant has been proposed to be characterized by prominent impairments in non-visual parietal functions (i.e., agraphia, alexia and apraxia), symptoms which are often present in posterior cortical atrophy (Mendez *et al*., 2002; McMonagle *et al*., 2006; Crutch *et al*., 2017). In the present study, we were unable to discern clear associations between any of the atrophy factors and dominant-parietal functions. Furthermore, outlining the impairments across cognitive domains revealed that none of the participants had a clearly isolated non-visual dominant parietal impairment, and the case we selected also had severe impairments on other domains and a low MMSE. The discrepancy between our findings and earlier studies, which formed the basis for the hypothesized dominant parietal variant, may again be that these were based on small studies or single case studies selected based on this particular phenotype. Also, these previous studies primarily focused on apraxia (which was not assessed in the present study) (De Renzi, 1986; Green *et al*., 1995; Aharon-Peretz *et al*., 1999). Another possible explanation for why we did not observe patients with isolated impairments in non-visual dominant parietal functions is that these individuals might have been less likely to be included because the clinical criteria for posterior cortical atrophy rely primarily on prominent visual features (Mendez *et al*., 2002; Tsai *et al*., 2011; Crutch *et al*., 2017).

We did detect atrophy factors that might be related to the caudal variant of posterior cortical atrophy, characterized by primary visual processing function deficits, namely the left and right-ventral factors. These factors encompassed the occipital regions proposed to be associated with the caudal variant. However, these factors also included inferior temporal and inferior parietal regions, so we were unable to discern a clearly caudal factor associated with primary visual processing in the main model (*K* = 4). When allowing for five factors (*K* = 5), we did observe a right-caudal factor (Supplemental Fig. 7), which was related to several primary visual processing tests, although these associations were not consistent across all tests (Supplemental Fig. 7). Remarkably, we also observed that individuals with high expression of the right-caudal factor tended to have high MMSE scores (Supplemental Fig. 8A) and there was a positive association between right-caudal factor expression and MMSE (Supplemental Fig. 8B), suggesting that individuals with prominent right-caudal atrophy tend to be in the earlier phases of clinical disease progression. These findings are in line with earlier observations that posterior cortical atrophy starts in the most posteriorly located regions of the brain and then spreads to lateral parieto-temporal cortices (Kennedy *et al*., 2012; Agosta *et al*., 2018; Firth *et al*., 2019), following a posterior-to-anterior pathway in line with the network-based degeneration hypothesis (Seeley *et al*., 2009). It may therefore be that descriptions of isolated impairments on primary visual processing and caudally located atrophy are based on individuals in an early stage of the disease, which would be in line with the observation that all posterior cortical atrophy participants show impairments in at least one primary visual domain (Lehmann *et al*., 2011*a*). Taken together, this suggests that the caudal variant may represent an early disease stage-related phenomenon, rather than a distinct variant of posterior cortical atrophy.

In summary, the aggregate of the literature in conjunction with the present study indicates that classification of individuals with posterior cortical atrophy into discrete variants is not straightforward. Our case series revealed that extreme phenotypes can indeed be detected among posterior cortical atrophy participants, and it is possible that these cases are the ones that have been described in the investigations that formed the basis for the described variants. However, the low prevalence of these extremes in combination with the considerable overlap in clinical and radiological characteristics among the rest of the population suggests that classification into discrete variants may have limited clinical value.

### Strengths and limitations

The main strengths of the present study include the relatively large, multi-center sample of extensively phenotyped posterior cortical atrophy participants. Furthermore, our data-driven approach allowed atrophy factors to be partly overlapping instead of completely distinct and allowed participants to express each atrophy factor to a certain degree. These characteristics make this approach more biologically plausible than *a priori* categorization of participants into mutually exclusive subgroups or selection of regions-of-interest to investigate. Furthermore, we excluded participants with negative amyloid-β biomarkers in order to increase the likelihood that individuals had posterior cortical atrophy due to Alzheimer’s disease (Renner *et al*., 2004; Tang-Wai *et al*., 2004; Montembeault *et al*., 2018), thereby excluding possible confounding effects of differences in underlying pathology as much as possible. However, this also constitutes a possible limitation as it has been shown that individuals with posterior cortical atrophy due to non-Alzheimer’s disease pathology show a different pattern of neurodegeneration (Montembeault *et al*., 2018). By focusing on participants with underlying Alzheimer’s disease pathology, we were not able to assess whether non-Alzheimer’s disease pathology could have formed the basis for the hypothesized phenotypical variants of posterior cortical atrophy. Another possible limitation of the present study is the retrospective inclusion of participants assessed from 2000 to 2017, which resulted in participants being selected based on different clinical criteria (Mendez *et al*., 2002; Tang-Wai *et al*., 2004; Crutch *et al*., 2017). However, there were no associations between date of inclusion and atrophy factor expression (Supplemental Fig. 9). Furthermore, our sample, while relatively large for a posterior cortical atrophy cohort, was relatively small (N=119), and we performed many comparisons. After correction for multiple comparisons, several of the associations between factor expressions and neuropsychological tests lost statistical significance. However, with these assessments we wanted to provide an overview of the associations between atrophy factors and cognitive impairments within the spectrum of posterior cortical atrophy, and we have included effect sizes in the results to allow the reader to draw her own conclusions.

### Conclusions and future directions

Akin to classifying Alzheimer’s disease patients into atypical variants (e.g., logopenic variant primary progressive aphasia or the dysexecutive/behavioral variant (Gorno-Tempini *et al*., 2011, Ossenkoppele *et al*., 2015*c*)), subtyping posterior cortical atrophy into variants suffers from arbitrary criteria that constrain the wealth of clinical and radiological variability into categorical entities. These classifications are mostly useful in a clinical setting, to aid in an early and accurate diagnosis and to direct patient care as well as aiding in selection for clinical trials (Scheltens *et al*., 2017). However, when only the extremes of an already relatively rare syndrome are captured by this classification, the clinical utility becomes limited and, for clinical purposes, categorizing posterior cortical atrophy as a single entity might be sufficiently specific. Elucidating the link between clinical heterogeneity and neurobiological differences may, however, be useful in a research setting to assess mechanisms leading to selective vulnerability in neurodegenerative diseases (Mattsson *et al*., 2016). Posterior cortical atrophy may be the easiest to distinguish from “typical” Alzheimer’s disease in the early stages of the disease (Lehmann *et al*., 2012; Firth *et al*., 2019) and hypothetical models of Alzheimer’s disease suggest that tau aggregation and hypometabolism precede neurodegeneration (Jack and Holtzman, 2013). This indicates that successfully identifying phenotypical variants of posterior cortical atrophy may rely on early detection using, for example, tau-PET or FDG-PET, which have already been shown to distinguish posterior cortical atrophy from typical Alzheimer’s disease (Nestor *et al*., 2003; Whitwell *et al*., 2007; Rosenbloom *et al*., 2011; Ossenkoppele *et al*., 2012, 2015*d*; Day *et al*., 2017; Panegyres *et al*., 2017). Another possible avenue to detect phenotypical variants of posterior cortical atrophy may be the assessment of functional connectivity (Lehmann *et al*., 2015; Agosta *et al*., 2018; Fredericks *et al*., 2019), as emerging evidence points to the spread of neurodegeneration along intrinsic functional brain networks. Aside from neuroimaging factors related to regional vulnerability, it has also been shown that there are genetic risk factors that convey a specific risk to posterior cortical atrophy (Schott *et al*., 2016). In addition, a recent study has found that mathematical and visuospatial learning difficulties are related to visuospatial predominant clinical syndromes, which indicates that neurodevelopment might also be related to vulnerability of specific brain networks that predisposes an individual to show network failure in these systems when neurodegenerative diseases emerge in later-life (Miller *et al*., 2018).

These emerging findings help to elucidate the intricate pathways that eventually result in discrete clinical syndromes and indicate that regional susceptibility to pathology is most likely multifactorial. Considering the interplay between different susceptibility factors, future examinations assessing regional vulnerability will therefore require multi-modal assessment with large sample sizes. Owing to the relatively low prevalence of posterior cortical atrophy, obtaining sufficient cohorts exclusively containing individuals with posterior cortical atrophy will remain challenging. For now, it might be prudent to focus on the entire Alzheimer’s disease spectrum and examine factors related to particular vulnerability for developing posterior cortical atrophy rather than specific variations of this already relatively rare syndrome.

## Acknowledgements

We thank Sarah Remmers for her comments on the neuropsychological data, JQ Alida Chen for helping to design the figures and Paul K Crane for proof-reading the manuscript.

Research of the Alzheimer Center Amsterdam is part of the neurodegeneration research program of the Neuroscience Campus Amsterdam. The Alzheimer Center Amsterdam is supported by Alzheimer Nederland and Stichting Amsterdam UMC Fonds. The clinical database structure was developed with funding from Stichting Dioraphte. Wiesje M van der Flier holds the Pasman chair.

## References

Agosta F, Mandic-Stojmenovic G, Canu E, Stojkovic T, Imperiale F, Caso F, et al. Functional and structural brain networks in posterior cortical atrophy: A two-centre multiparametric MRI study. NeuroImage Clin 2018; 19: 901–910.

Aharon-Peretz J, Israel O, Goldsher D, Peretz A. Posterior Cortical Atrophy Variants of Alzheimer’s Disease. Dement Geriatr Cogn Disord 1999; 10: 483–487.

Ahmed S, Loane C, Bartels S, Zamboni G, Mackay C, Baker I, et al. Lateral parietal contributions to memory impairment in posterior cortical atrophy. NeuroImage Clin 2018; 20: 252–259.

Alves J, Soares JM, Sampaio A, Gonçalves ÓF. Posterior cortical atrophy and Alzheimer’s disease: a meta-analytic review of neuropsychological and brain morphometry studies. Brain Imaging Behav 2013; 7: 353–361.

Ashburner J. A fast diffeomorphic image registration algorithm. Neuroimage 2007; 38: 95–113.

Benson DF, Davis RJ, Snyder BD. Posterior cortical atrophy. Arch Neurol 1988; 45: 789–93.

Borruat F-X. Posterior Cortical Atrophy: Review of the Recent Literature. Curr Neurol Neurosci Rep 2013; 13: 406.

Boyd CD, Tierney M, Wassermann EM, Spina S, Oblak AL, Ghetti B, et al. Visuoperception test predicts pathologic diagnosis of Alzheimer disease in corticobasal syndrome. Neurology 2014; 83: 510–519.

Chan D, Crutch SJ, Warrington EK. A disorder of colour perception associated with abnormal colour after-images: a defect of the primary visual cortex. J Neurol Neurosurg Psychiatry 2001; 71: 515–7.

Crutch SJ, Lehmann M, Schott JM, Rabinovici GD, Rossor MN, Fox NC. Posterior cortical atrophy. Lancet Neurol 2012; 11: 170–178.

Crutch SJ, Lehmann M, Warren JD, Rohrer JD. The language profile of posterior cortical atrophy. J Neurol Neurosurg Psychiatry 2013; 84: 460–466.

Crutch SJ, Schott JM, Rabinovici GD, Murray M, Snowden JS, van der Flier WM, et al. Consensus classification of posterior cortical atrophy. Alzheimer’s Dement 2017; 13: 870–884.

Day GS, Gordon BA, Jackson K, Christensen JJ, Rosana Ponisio M, Su Y, et al. Tau-PET Binding Distinguishes Patients With Early-stage Posterior Cortical Atrophy From Amnestic Alzheimer Disease Dementia. Alzheimer Dis Assoc Disord 2017; 31: 87–93.

Dubois B, Feldman HH, Jacova C, Dekosky ST, Barberger-Gateau P, Cummings J, et al. Research criteria for the diagnosis of Alzheimer’s disease: revising the NINCDS-ADRDA criteria. Lancet Neurol 2007; 6: 734–46.

Firth NC, Primativo S, Marinescu R-V, Shakespeare TJ, Suarez-Gonzalez A, Lehmann M, et al. Longitudinal neuroanatomical and cognitive progression of posterior cortical atrophy. Brain 2019.

van der Flier WM, Scheltens P. Amsterdam Dementia Cohort: Performing Research to Optimize Care. J Alzheimers Dis 2018; 62: 1091–1111.

Fredericks CA, Brown JA, Deng J, Kramer A, Ossenkoppele R, Rankin K, et al. Intrinsic connectivity networks in posterior cortical atrophy: A role for the pulvinar? NeuroImage Clin 2019; 21: 101628.

Galton CJ, Patterson K, Xuereb JH, Hodges JR. Atypical and typical presentations of Alzheimer’s disease: a clinical, neuropsychological, neuroimaging and pathological study of 13 cases. Brain 2000; 123 Pt 3: 484–98.

Gorno-Tempini ML, Hillis AE, Weintraub S, Kertesz A, Mendez M, Cappa SF, et al. Classification of primary progressive aphasia and its variants. Neurology 2011; 76: 1006–14.

Green RC, Goldstein FC, Mirra SS, Alazraki NP, Baxt JL, Bakay RA. Slowly progressive apraxia in Alzheimer’s disease. J Neurol Neurosurg Psychiatry 1995; 59: 312–5.

Groot C, Van Loenhoud ACAC, Barkhof F, van Berckel BNMBNM, Koene T, Teunissen CCCC, et al. Differential effects of cognitive reserve and brain reserve on cognition in Alzheimer disease. Neurology 2018; 90: e149–e156.

Grossi D, Soricelli A, Ponari M, Salvatore E, Quarantelli M, Prinster A, et al. Structural connectivity in a single case of progressive prosopagnosia: The role of the right inferior longitudinal fasciculus. Cortex 2014; 56: 111–120.

Huberle E, Rupek P, Lappe M, Karnath H-O. Perception of biological motion in visual agnosia. Front Behav Neurosci 2012; 6: 56.

Jack CR, Holtzman DM. Biomarker modeling of Alzheimer’s disease. Neuron 2013; 80: 1347–1358.

La Joie R, Perrotin A, Barré L, Hommet C, Mézenge F, Ibazizene M, et al. Region-specific hierarchy between atrophy, hypometabolism, and β-amyloid (Aβ) load in Alzheimer’s disease dementia. J Neurosci 2012; 32: 16265–73.

Kaeser P-F, Ghika J, Borruat F-X. Visual signs and symptoms in patients with the visual variant of Alzheimer disease. BMC Ophthalmol 2015; 15: 65.

ten Kate M, Barkhof F, Visser PJ, Teunissen CE, Scheltens P, van der Flier WM, et al. Amyloid-independent atrophy patterns predict time to progression to dementia in mild cognitive impairment. Alzheimers Res Ther 2017; 9: 73.

Kennedy J, Lehmann M, Sokolska MJ, Archer H, Warrington EK, Fox NC, et al. Visualizing the emergence of posterior cortical atrophy. Neurocase 2012; 18: 248–57.

Koedam ELGE, Lehmann M, van der Flier WM, Scheltens P, Pijnenburg YAL, Fox N, et al. Visual assessment of posterior atrophy development of a MRI rating scale. Eur Radiol 2011; 21: 2618–2625.

Kramer JH, Jurik J, Sha SJ, Rankin KP, Rosen HJ, Johnson JK, et al. Distinctive neuropsychological patterns in frontotemporal dementia, semantic dementia, and Alzheimer disease. Cogn Behav Neurol 2003; 16: 211–8.

Lehmann M, Barnes J, Ridgway GR, Ryan NS, Warrington EK, Crutch SJ, et al. Global gray matter changes in posterior cortical atrophy: a serial imaging study. Alzheimers Dement 2012; 8: 502–12.

Lehmann M, Barnes J, Ridgway GR, Wattam-Bell J, Warrington EK, Fox NC, et al. Basic Visual Function and Cortical Thickness Patterns in Posterior Cortical Atrophy. Cereb Cortex 2011; 21: 2122–2132.

Lehmann M, Crutch SJ, Ridgway GR, Ridha BH, Barnes J, Warrington EK, et al. Cortical thickness and voxel-based morphometry in posterior cortical atrophy and typical Alzheimer’s disease. Neurobiol Aging 2011; 32: 1466–1476.

Lehmann M, Ghosh PM, Madison C, Laforce R, Corbetta-Rastelli C, Weiner MW, et al. Diverging patterns of amyloid deposition and hypometabolism in clinical variants of probable Alzheimer’s disease. Brain 2013; 136: 844–858.

Lehmann M, Madison C, Ghosh PM, Miller ZA, Greicius MD, Kramer JH, et al. Loss of functional connectivity is greater outside the default mode network in nonfamilial early-onset Alzheimer’s disease variants. Neurobiol Aging 2015; 36: 2678–2686.

Levine DN, Lee JM, Fisher CM. The visual variant of Alzheimer’s disease: A clinicopathologic case study. Neurology 1993; 43: 305–305.

van Loenhoud AC, Wink AM, Groot C, Verfaillie SCJ, Twisk J, Barkhof F, et al. A neuroimaging approach to capture cognitive reserve: Application to Alzheimer’s disease. Hum Brain Mapp 2017; 38

Magnin E, Sylvestre G, Lenoir F, Dariel E, Bonnet L, Chopard G, et al. Logopenic syndrome in posterior cortical atrophy. J Neurol 2013; 260: 528–533.

Marinescu R V., Eshaghi A, Lorenzi M, Young AL, Oxtoby NP, Garbarino S, et al. DIVE: A spatiotemporal progression model of brain pathology in neurodegenerative disorders. Neuroimage 2019; 192: 166–177.

Mattsson N, Schott JM, Hardy J, Turner MR, Zetterberg H. Selective vulnerability in neurodegeneration: insights from clinical variants of Alzheimer’s disease. J Neurol Neurosurg Psychiatry 2016; 87: 1000–1004.

McMonagle P, Deering F, Berliner Y, Kertesz A. The cognitive profile of posterior cortical atrophy. Neurology 2006; 66: 331–338.

Mendez MF, Ghajarania M, Perryman KM. Posterior Cortical Atrophy: Clinical Characteristics and Differences Compared to Alzheimer’s Disease. Dement Geriatr Cogn Disord 2002; 14: 33–40.

Migliaccio R, Agosta F, Rascovsky K, Karydas A, Bonasera S, Rabinovici GD, et al. Clinical syndromes associated with posterior atrophy: Early age at onset AD spectrum. Neurology 2009; 73: 1571–1578.

Migliaccio R, Agosta F, Scola E, Magnani G, Cappa SF, Pagani E, et al. Ventral and dorsal visual streams in posterior cortical atrophy: A DT MRI study. Neurobiol Aging 2012; 33: 2572–2584.

Miller ZA, Rosenberg L, Santos-Santos MA, Stephens M, Allen IE, Hubbard HI, et al. Prevalence of Mathematical and Visuospatial Learning Disabilities in Patients With Posterior Cortical Atrophy. JAMA Neurol 2018; 75: 728.

Montembeault M, Brambati SM, Lamari F, Michon A, Samri D, Epelbaum S, et al. Atrophy, metabolism and cognition in the posterior cortical atrophy spectrum based on Alzheimer’s disease cerebrospinal fluid biomarkers. NeuroImage Clin 2018; 20: 1018–1025.

Nestor PJ, Caine D, Fryer TD, Clarke J, Hodges JR. The topography of metabolic deficits in posterior cortical atrophy (the visual variant of Alzheimer’s disease) with FDG-PET. J Neurol Neurosurg Psychiatry 2003; 74: 1521–9.

Ossenkoppele R, Cohn-Sheehy BI, La Joie R, Vogel JW, Möller C, Lehmann M, et al. Atrophy patterns in early clinical stages across distinct phenotypes of Alzheimer’s disease. Hum Brain Mapp 2015; 36: 4421–4437.

Ossenkoppele R, Mattsson N, Teunissen CE, Barkhof F, Pijnenburg Y, Scheltens P, et al. Cerebrospinal fluid biomarkers and cerebral atrophy in distinct clinical variants of probable Alzheimer’s disease. Neurobiol Aging 2015; 36: 2340–2347.

Ossenkoppele R, Pijnenburg YAL, Perry DC, Cohn-Sheehy BI, Scheltens NME, Vogel JW, et al. The behavioural/dysexecutive variant of Alzheimer’s disease: clinical, neuroimaging and pathological features. Brain 2015; 138: 2732–49.

Ossenkoppele R, Prins ND, Pijnenburg YAL, Lemstra AW, van der Flier WM, Adriaanse SF, et al. Impact of molecular imaging on the diagnostic process in a memory clinic. Alzheimer’s Dement 2013; 9: 414–421.

Ossenkoppele R, Schonhaut DR, Baker SL, James P, Neil O, Janabi M, et al. Tau, Amyloid, and Hypometabolism in a Patient with Posterior Cortical Atrophy. 2015; 77: 338–342.

Ossenkoppele R, Zwan MD, Tolboom N, van Assema DME, Adriaanse SF, Kloet RW, et al. Amyloid burden and metabolic function in early-onset Alzheimer’s disease: parietal lobe involvement. Brain 2012; 135: 2115–25.

Panegyres PK, Goh J, McCarthy M, Campbell AI. The Nature and Natural History of Posterior Cortical Atrophy Syndrome. Alzheimer Dis Assoc Disord 2017; 31: 295–306.

Phillips JS, Da Re F, Irwin DJ, McMillan CT, Vaishnavi SN, Xie SX, et al. Longitudinal progression of grey matter atrophy in non-amnestic Alzheimer’s disease. Brain 2019; 142: 1701–1722.

Rabinovici GD, Furst AJ, Alkalay A, Racine CA, O’Neil JP, Janabi M, et al. Increased metabolic vulnerability in early-onset Alzheimer’s disease is not related to amyloid burden. Brain 2010; 133: 512–528.

Renner JA, Burns JM, Hou CE, McKeel DW, Storandt M, Morris JC. Progressive posterior cortical dysfunction: a clinicopathologic series. Neurology 2004; 63: 1175–80.

De Renzi E. Slowly progressive visual agnosia or apraxia without dementia. Cortex 1986; 22: 171–80.

Rosenbloom MH, Alkalay A, Agarwal N, Baker SL, O’Neil JP, Janabi M, et al. Distinct clinical and metabolic deficits in PCA and AD are not related to amyloid distribution. Neurology 2011; 76: 1789–96.

Ross SJ, Graham N, Stuart-Green L, Prins M, Xuereb J, Patterson K, et al. Progressive biparietal atrophy: an atypical presentation of Alzheimer’s disease. J Neurol Neurosurg Psychiatry 1996; 61: 388–95.

Scheltens NME, Tijms BM, Koene T, Barkhof F, Teunissen CE, Wolfsgruber S, et al. Cognitive subtypes of probable Alzheimer’s disease robustly identified in four cohorts. Alzheimers Dement 2017; 13: 1226–1236.

Schott JM, Crutch SJ. Posterior Cortical Atrophy. Contin Lifelong Learn Neurol 2019; 25: 52–75.

Schott JM, Crutch SJ, Carrasquillo MM, Uphill J, Shakespeare TJ, Ryan NS, et al. Genetic risk factors for the posterior cortical atrophy variant of Alzheimer’s disease. Alzheimer’s Dement 2016; 12: 862–871.

Seeley WW, Crawford RK, Zhou J, Miller BL, Greicius MD. Neurodegenerative Diseases Target Large-Scale Human Brain Networks. Neuron 2009; 62: 42–52.

Snowden JS, Stopford CL, Julien CL, Thompson JC, Davidson Y, Gibbons L, et al. Cognitive phenotypes in Alzheimer’s disease and genetic risk. Cortex 2007; 43: 835–45.

Tang-Wai DF, Graff-Radford NR, Boeve BF, Dickson DW, Parisi JE, Crook R, et al. Clinical, genetic, and neuropathologic characteristics of posterior cortical atrophy. Neurology 2004; 63: 1168–74.

Tijms BM, Willemse EAJ, Zwan MD, Mulder SD, Visser PJ, van Berckel BNM, et al. Unbiased Approach to Counteract Upward Drift in Cerebrospinal Fluid Amyloid-β 1–42 Analysis Results. Clin Chem 2018; 64: 576–585.

Tsai P-H, Teng E, Liu C, Mendez MF. Posterior Cortical Atrophy. Am J Alzheimer’s Dis Other Dementiasr 2011; 26: 413–418.

Ungerleider LG, Haxby J V. ‘What’ and ‘where’ in the human brain. Curr Opin Neurol 1994; 4: 157–165.

Whitwell JL, Jack CR, Kantarci K, Weigand SD, Boeve BF, Knopman DS, et al. Imaging correlates of posterior cortical atrophy. Neurobiol Aging 2007; 28: 1051–61.

Zhang X, Mormino EC, Sun N, Sperling RA, Sabuncu MR, Yeo BTT. Bayesian model reveals latent atrophy factors with dissociable cognitive trajectories in Alzheimer’s disease. Proc Natl Acad Sci 2016; 113: E6535–E6544.

Zwan M, van Harten A, Ossenkoppele R, Bouwman F, Teunissen C, Adriaanse S, et al. Concordance between cerebrospinal fluid biomarkers and [11C]PIB PET in a memory clinic cohort. J Alzheimers Dis 2014; 41: 801–7.

