## Supplement for "Data-driven detection of latent atrophy factors related to phenotypical variants of posterior cortical atrophy"

**Supplemental material**

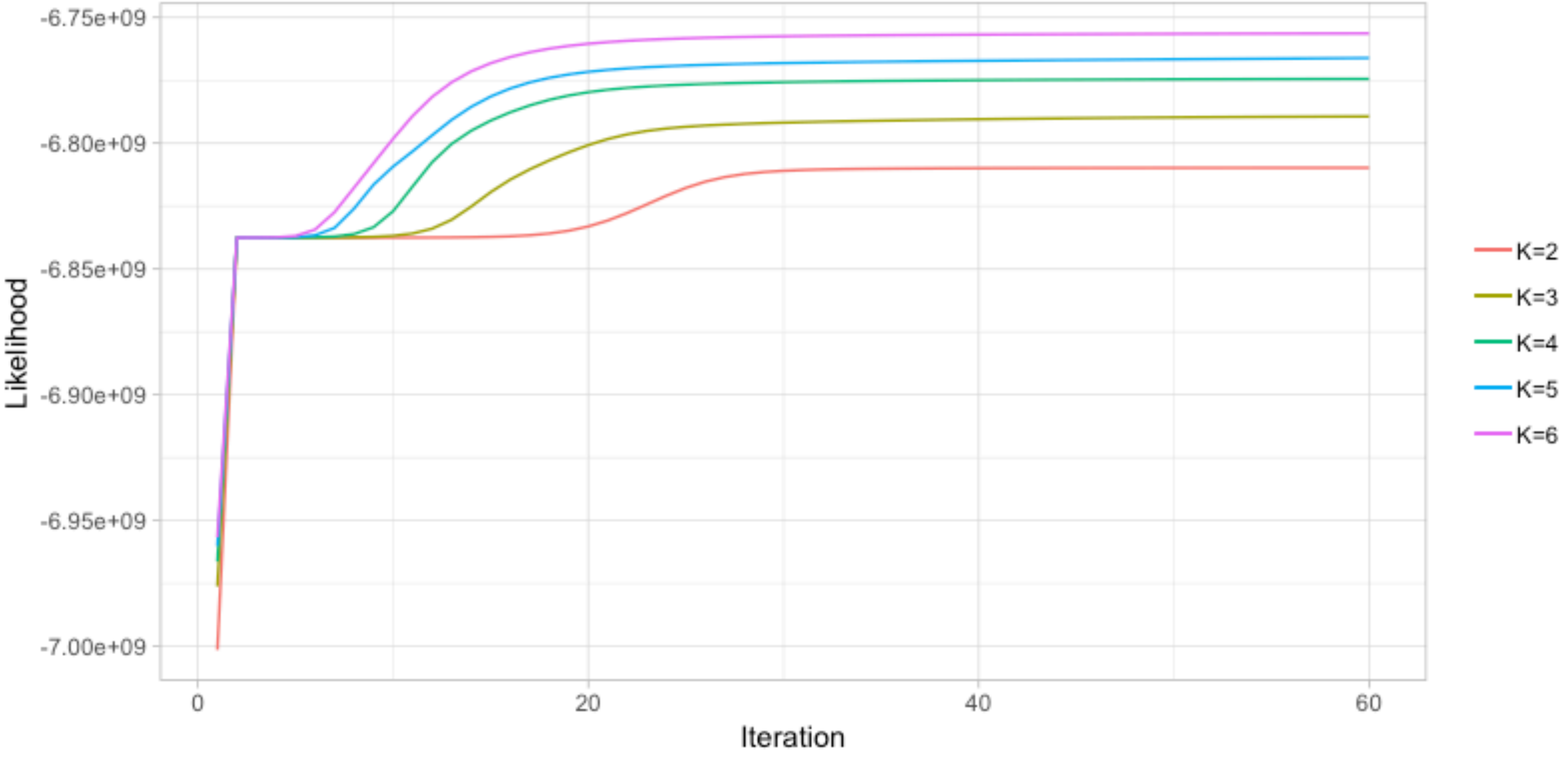

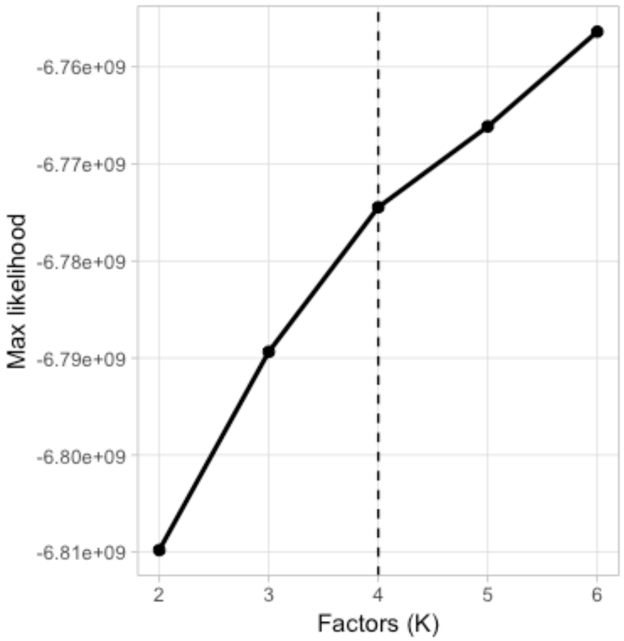


**Supplemental Figure 1. Likelihood convergence and max log likelihood comparisons across latent Dirichlet allocation models**

Panel A displays the likelihood conversion across iterations (max 60) for each of the latent Dirichlet allocation models, K=2 through 6, in the combined sample. Likelihood stabilizes around 30 to 40 iterations for all factors and increases with an increasing number of factor (K+1). Panel B displays the max likelihood (y-axis) for each of the models (x-axis). The dashed line represents the point where the increases in max likelihood starts to stabilize according to the “elbow method”. As expected, likelihood convergences revealed that model fit improved when increasing *K*, but the increase in max likelihood across models stabilized at $K = 4$.

### Amsterdam UMC
(N=69)

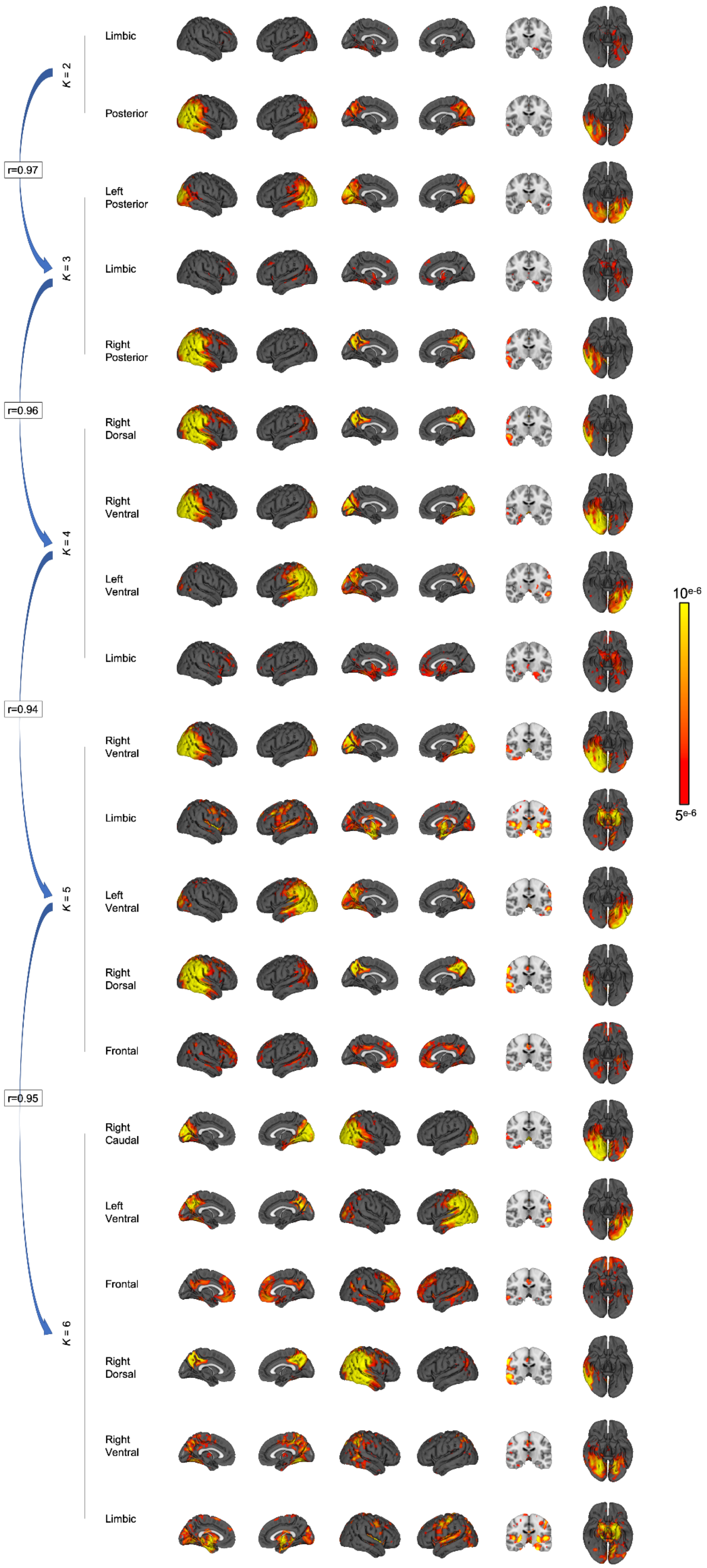

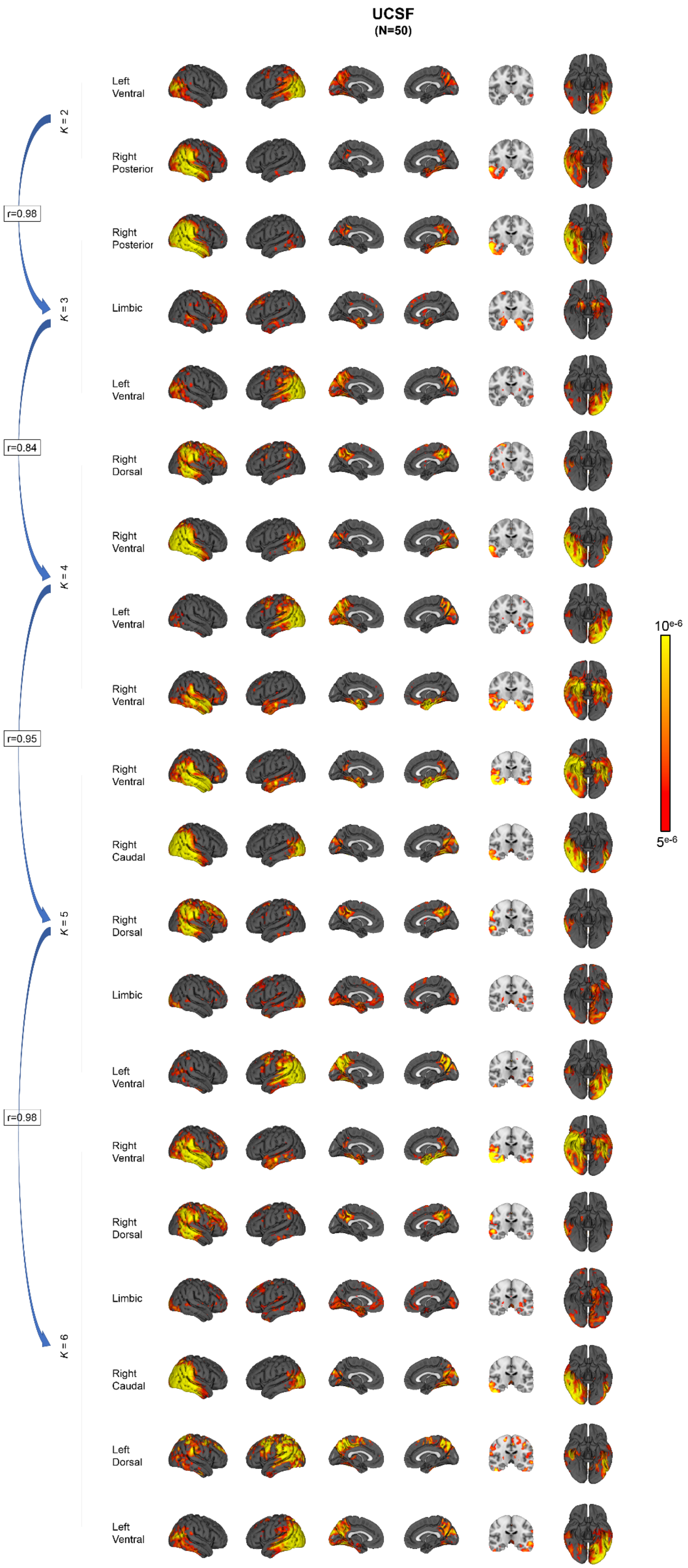

UCSF
(N=50)
K = 2
K = 3
K = 4
K = 5
K = 6
r=0.98
r=0.84
r=0.95
r=0.98
Left Ventral
Right Posterior
Right Posterior
Limbic
Left Ventral
Right Dorsal
Right Ventral
Left Ventral
Right Ventral
Right Ventral
Right Caudal
Right Dorsal
Limbic
Left Ventral
Right Ventral
Right Dorsal
Limbic
Right Caudal
Left Dorsal
Left Ventral
10e-6
5e-6

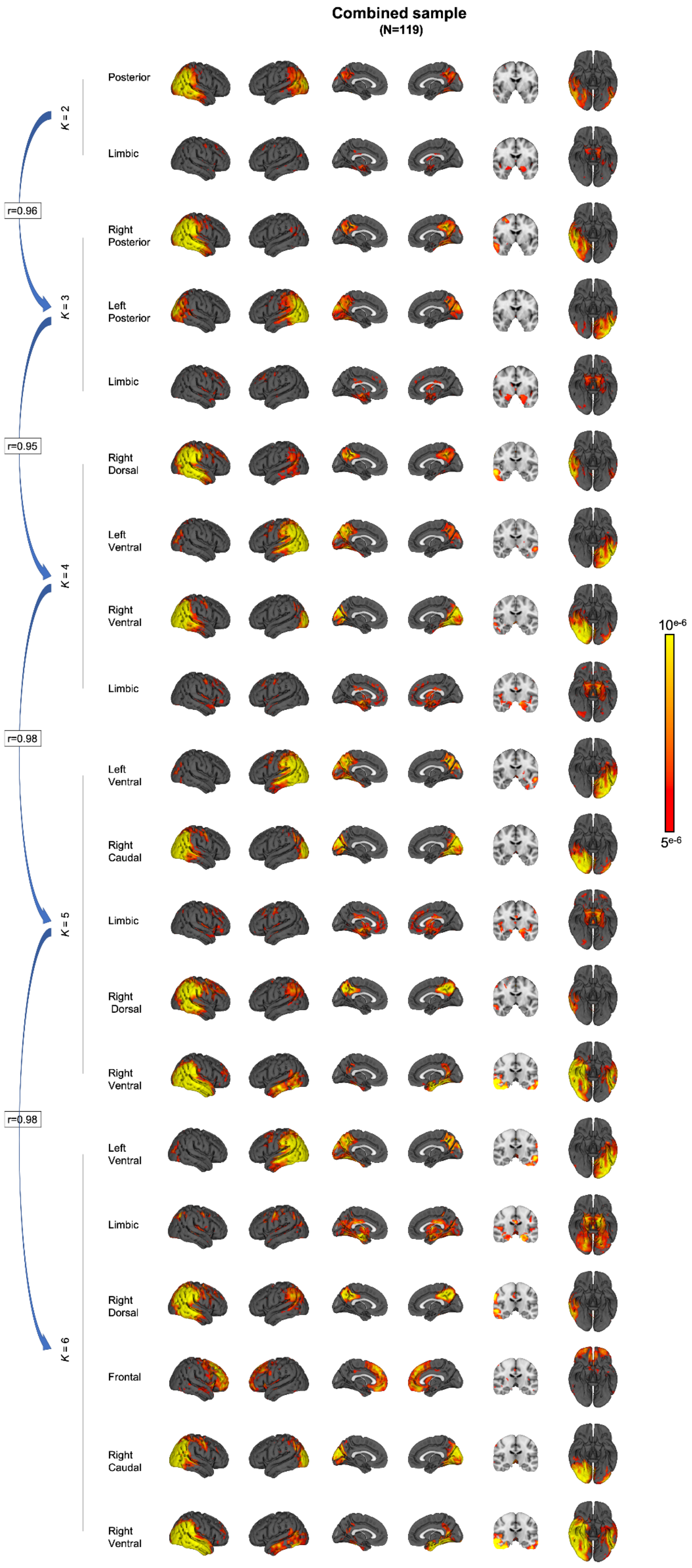

Combined sample
(N=119)
K = 2
K = 3
K = 4
K = 5
K = 6
r=0.96
r=0.95
r=0.98
r=0.98
Posterior
Limbic
Right Posterior
Left Posterior
Limbic
Right Dorsal
Left Ventral
Right Ventral
Limbic
Left Ventral
Right Caudal
Limbic
Right Dorsal
Right Ventral
Left Ventral
Limbic
Right Dorsal
Frontal
Right Caudal
Right Ventral
10e-6
5e-6

**Supplemental Figure 2. Atrophy factors revealed by latent Dirichlet allocation (*K*=2 through 6)**

Intensity of voxels signify the probability [Pr(Voxel | Factor)] of a voxel belonging to one of the factors. Figure 1A displays factors for the AUMC cohort. 1B for the UCSF cohort and 1C for the combined cohort. Scale is truncated at [Pr(Voxel | Factor)] = $5^{e-6}$ for visualization purposes. We determined how factor estimations from $K$ = 2 to 6 factors were related by performing an exhaustive search to quantify the possibility that two atrophy patterns in the ($K$+1) factor model were subdivisions of a pattern in the $K$-factor model. Using the Hungarian matching algorithm, the correlation of Pr(Voxel | Factor) between corresponding pairs of factors was obtained and averaged across all pairs of factors, resulting in an average correlation value indicating the quality of the split. The average correlation of Pr(Voxel | Factor) for factors 2 to 6 in the combined sample (ranging from $r$=0.95-0.98) indicates that the latent atrophy factors are ordered in a hierarchical fashion. Panel A displays results for the Amsterdam UMC sample, panel B for the UCSF sample and panel C for the combined sample.

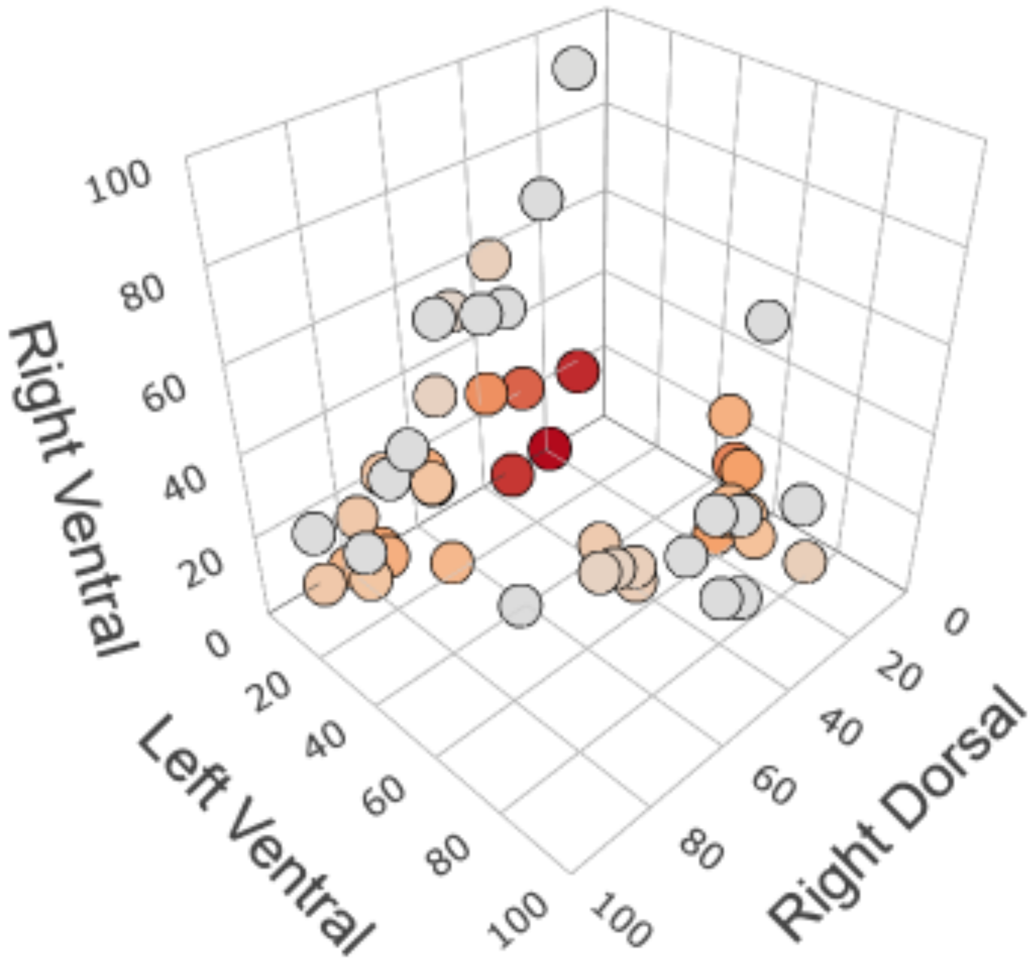

Right Ventral
100
80
60
40
20
0
Left Ventral
20
40
60
80
100
Right Dorsal
0
20
40
60
80
100

**Supplemental Figure 3. Atrophy factors compositions for the separate samples**

This 4D plots displays the factors left-ventral. right-dorsal and right-ventral on the x. y and z axes and the limbic factor is displayed by the color gradient of the markers. Expressions of the four factors adds up to 100%. Panel A displays factor compositions for the AUMC sample and panel B for the UCSF sample.

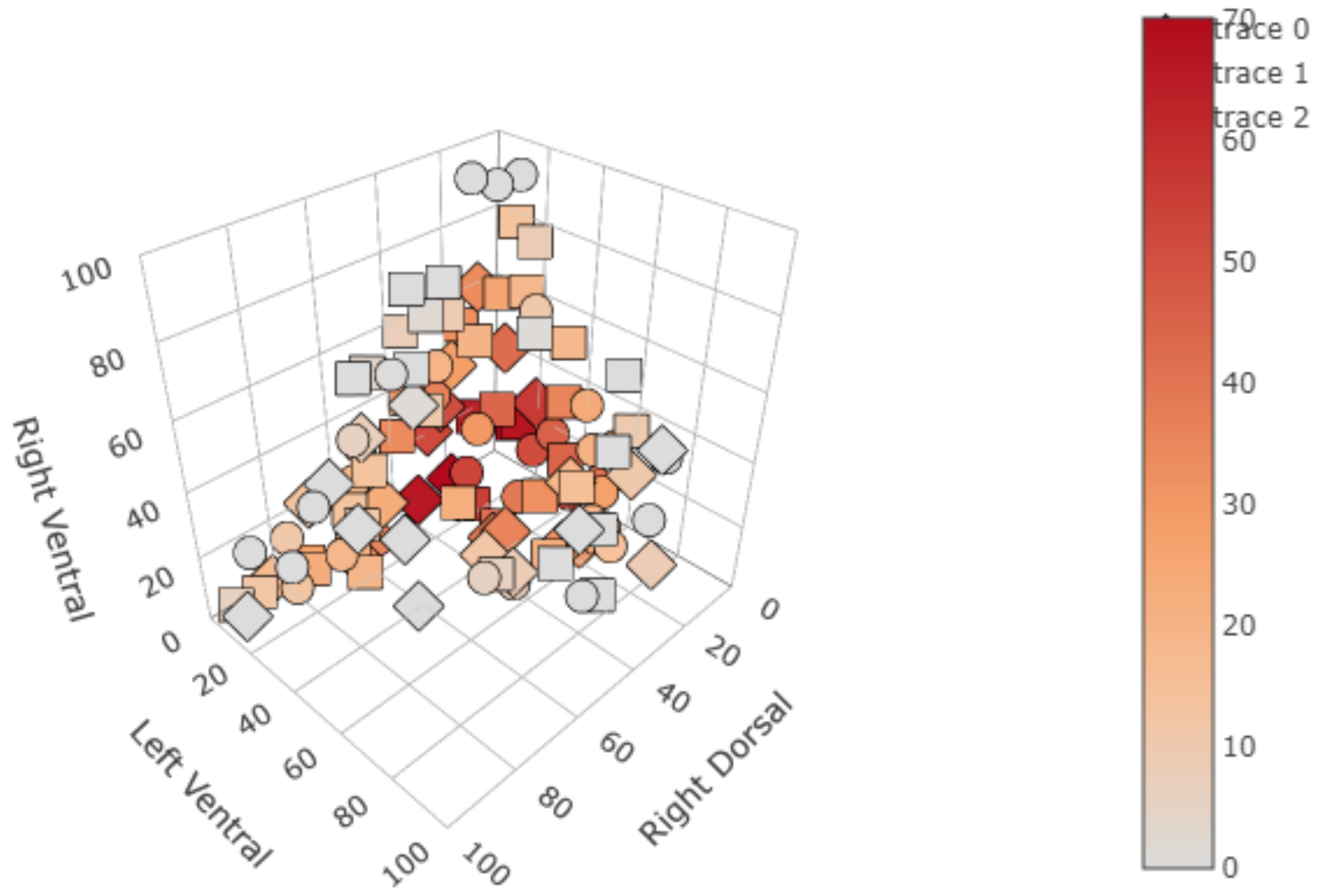

Right Ventral
Left Ventral
Right Dorsal
0
20
40
60
80
100
trace 0
trace 1
trace 2
70
60
50
40
30
20
10
0

**Supplemental Figure 4. Atrophy factors compositions for groups stratified according to clinical disease severity**

This 4D plots displays the factors left-ventral. right-dorsal and right-ventral on the x. y and z axes and the limbic factor is displayed by the color gradient of the markers. Expressions of the four factors adds up to 100%. The marker symbols represent the 3 clinical disease severity groups. Diamonds = MMSE 31-24, squares = MMSE 24-18 and circles = MMSE 18-6.

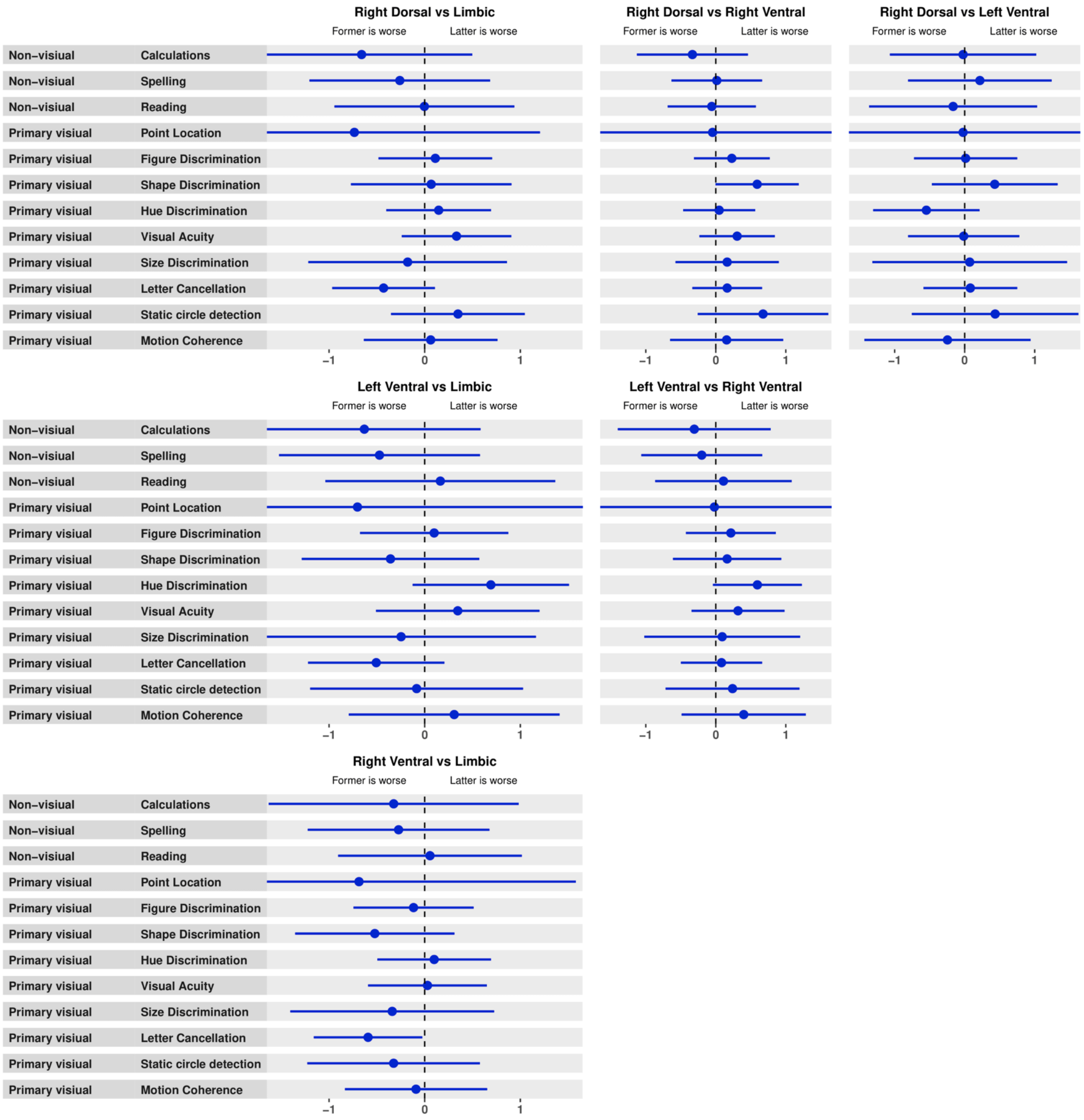
Right Dorsal vs Limbic
Right Dorsal vs Right Ventral
Right Dorsal vs Left Ventral
Left Ventral vs Limbic
Left Ventral vs Right Ventral
Right Ventral vs Limbic
Former is worse
Latter is worse
Non–visiual
Primary visiual
Calculations
Spelling
Reading
Point Location
Figure Discrimination
Shape Discrimination
Hue Discrimination
Visual Acuity
Size Discrimination
Letter Cancellation
Static circle detection
Motion Coherence
–1
0
1

**Supplemental Figure 5. Associations between factor expressions and neuropsychological tests assessing non-visual. dominant parietal and primary visual functions**

The plot contains relative cross-sectional effects from linear regression analyses. Lines indicate the 95% confidence intervals and a significant effect (uncorrected for multiple comparisons) is denoted by confidence intervals not including x=0. As in each model one of the factors is implicitly modelled. one of the factors serves as a reference to assess effects of all the others and each subplot displays results from the different factor comparisons

A

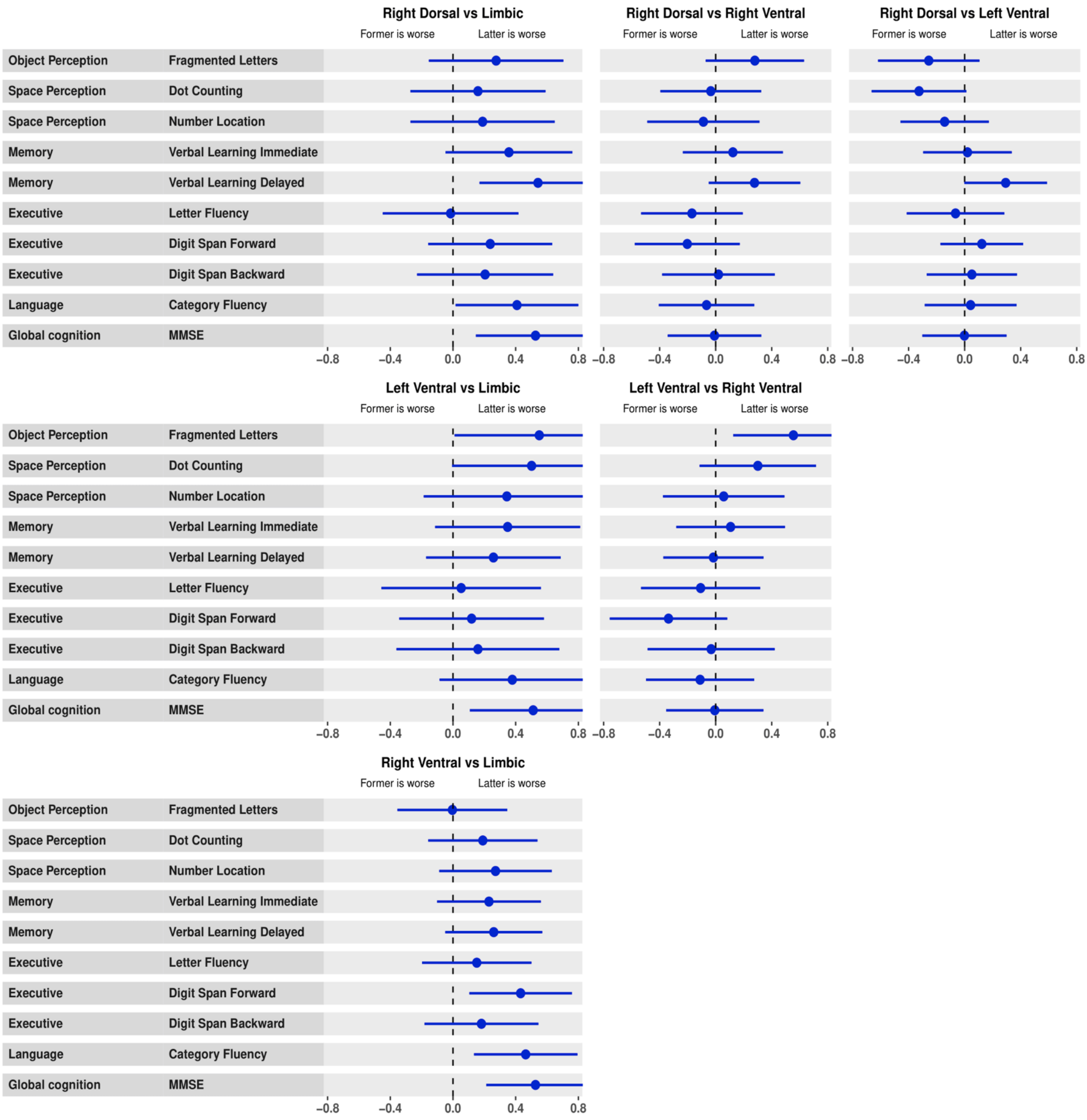

Right Dorsal vs Limbic
Right Dorsal vs Right Ventral
Right Dorsal vs Left Ventral
Left Ventral vs Limbic
Left Ventral vs Right Ventral
Right Ventral vs Limbic
Former is worse
Latter is worse
Object Perception
Fragmented Letters
Space Perception
Dot Counting
Number Location
Memory
Verbal Learning Immediate
Verbal Learning Delayed
Executive
Letter Fluency
Digit Span Forward
Digit Span Backward
Language
Category Fluency
Global cognition
MMSE
−0.8
−0.4
0.0
0.4
0.8

**B**

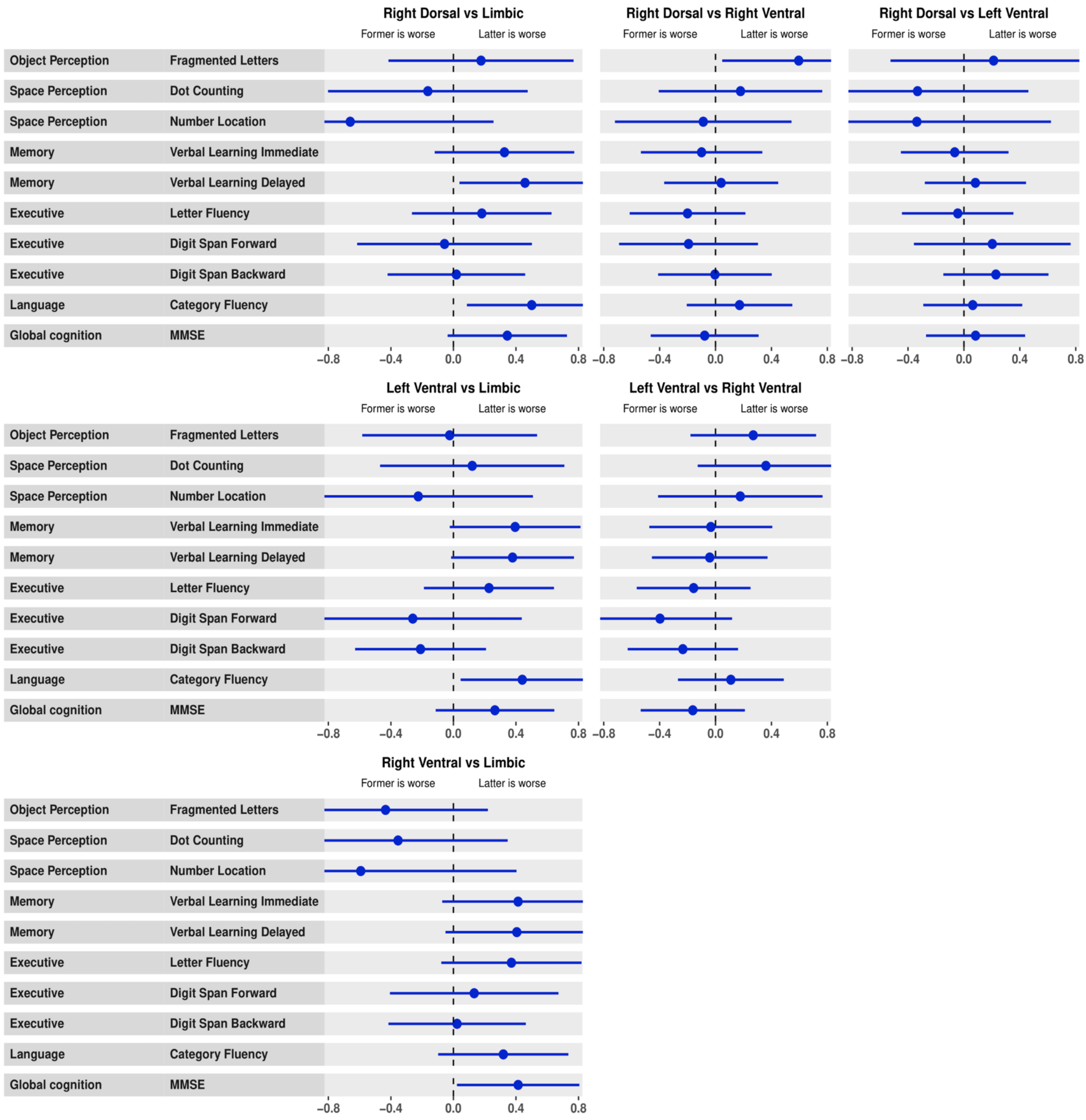
Right Dorsal vs Limbic
Right Dorsal vs Right Ventral
Right Dorsal vs Left Ventral
Left Ventral vs Limbic
Left Ventral vs Right Ventral
Right Ventral vs Limbic
Former is worse
Latter is worse
Object Perception
Fragmented Letters
Space Perception
Dot Counting
Number Location
Memory
Verbal Learning Immediate
Verbal Learning Delayed
Executive
Letter Fluency
Digit Span Forward
Digit Span Backward
Language
Category Fluency
Global cognition
MMSE
−0.8
−0.4
0.0
0.4
0.8

**Supplemental Figure 6. Associations between factor expressions and neuropsychological tests in the separate samples**

The plot contains relative cross-sectional effects from linear regression models. Lines indicate the 95% confidence intervals and a significant effect (uncorrected for multiple comparisons) is denoted by confidence intervals not including x=0. Panel A displays results for the Amsterdam UMC sample and panel B for the UCSF sample.

**MMSE**

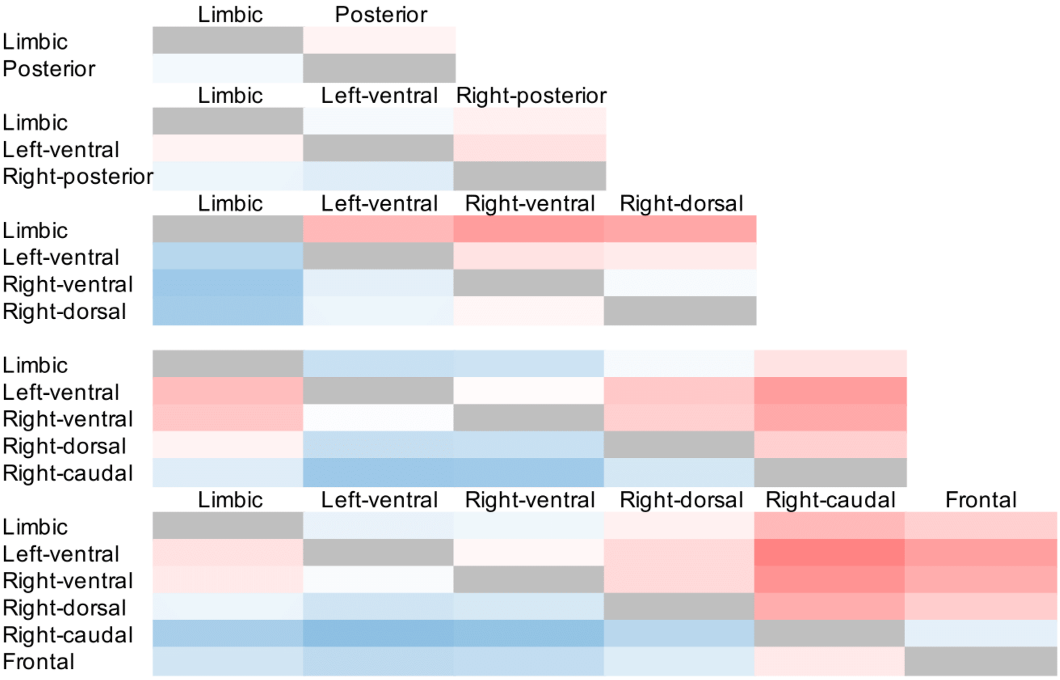


**Verbal Learning, immediate recall**

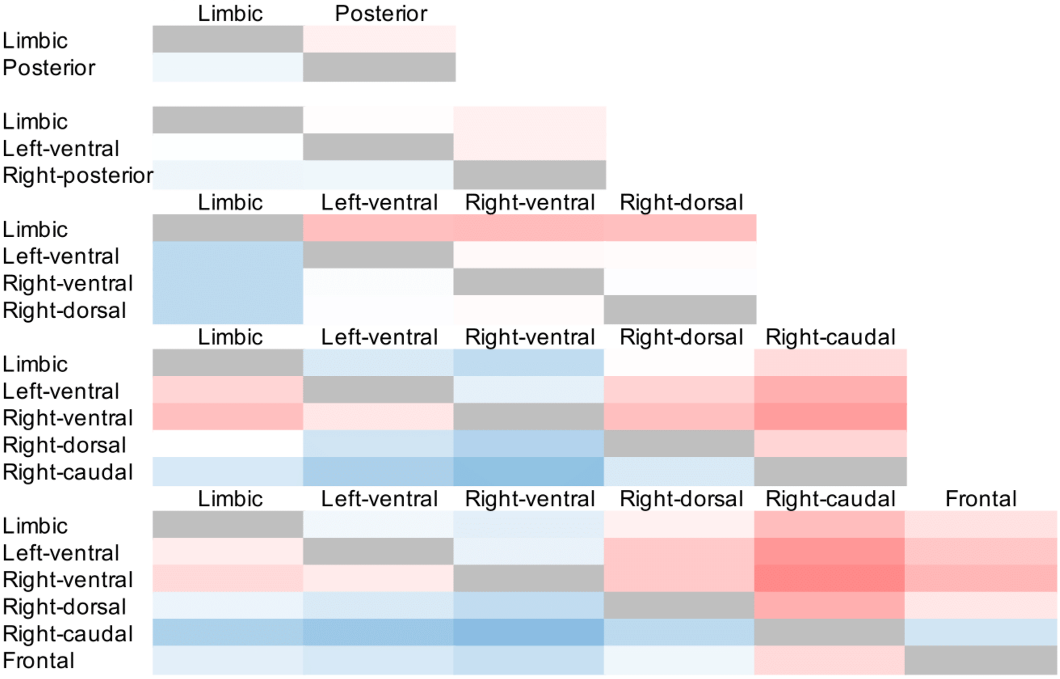


**Verbal Learning, delayed recall**

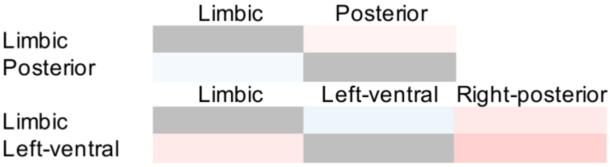

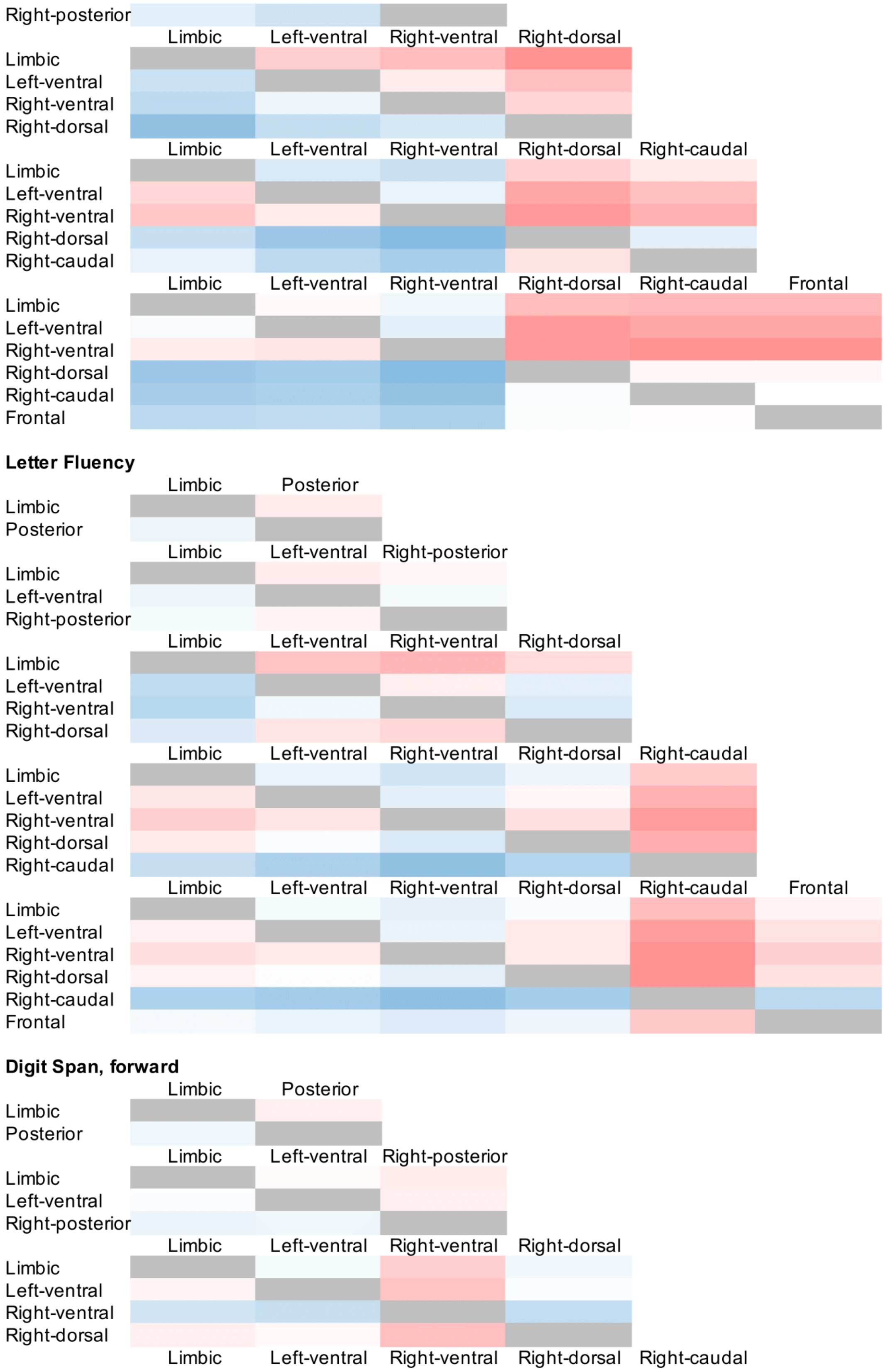
Right-posterior
Limbic Left-ventral Right-ventral Right-dorsal
Limbic
Left-ventral
Right-ventral
Right-dorsal
Limbic Left-ventral Right-ventral Right-dorsal Right-caudal
Limbic
Left-ventral
Right-ventral
Right-dorsal
Right-caudal
Limbic Left-ventral Right-ventral Right-dorsal Right-caudal Frontal
Limbic
Left-ventral
Right-ventral
Right-dorsal
Right-caudal
Frontal
Letter Fluency
Limbic Posterior
Limbic
Posterior
Limbic Left-ventral Right-posterior
Limbic
Left-ventral
Right-posterior
Limbic Left-ventral Right-ventral Right-dorsal
Limbic
Left-ventral
Right-ventral
Right-dorsal
Limbic Left-ventral Right-ventral Right-dorsal Right-caudal
Limbic
Left-ventral
Right-ventral
Right-dorsal
Right-caudal
Limbic Left-ventral Right-ventral Right-dorsal Right-caudal Frontal
Limbic
Left-ventral
Right-ventral
Right-dorsal
Right-caudal
Frontal
Digit Span, forward
Limbic Posterior
Limbic
Posterior
Limbic Left-ventral Right-posterior
Limbic
Left-ventral
Right-posterior
Limbic Left-ventral Right-ventral Right-dorsal
Limbic
Left-ventral
Right-ventral
Right-dorsal
Limbic Left-ventral Right-ventral Right-dorsal Right-caudal

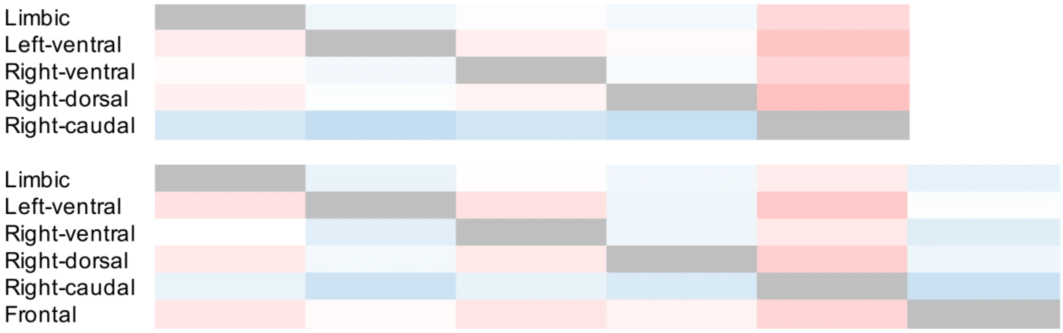


**Digit Span, backward**

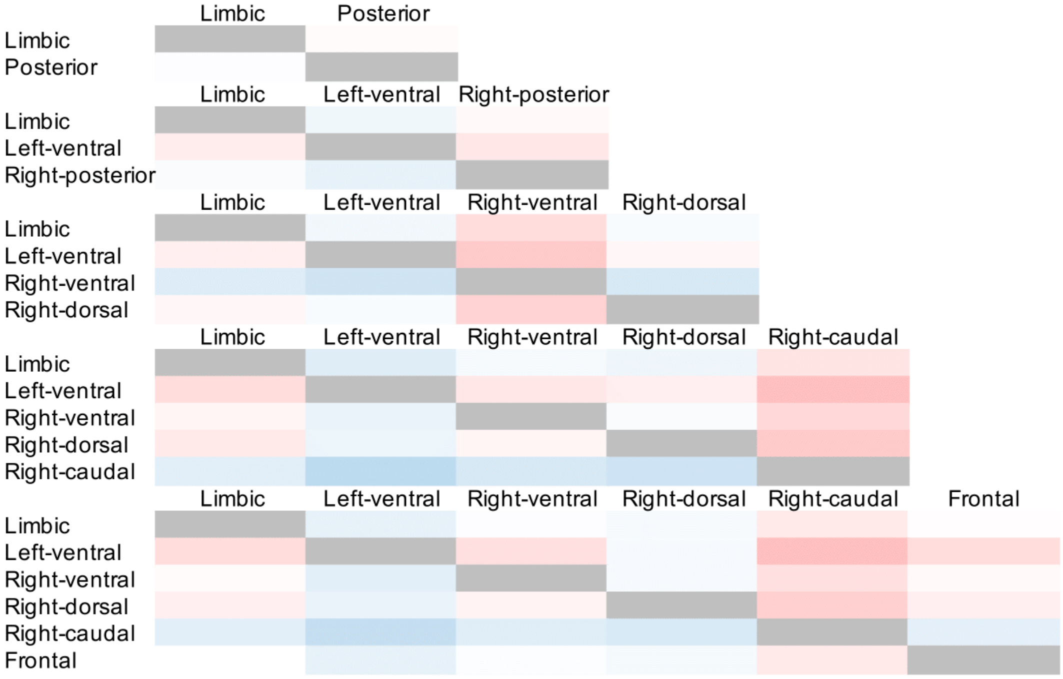


**Category Fluency**

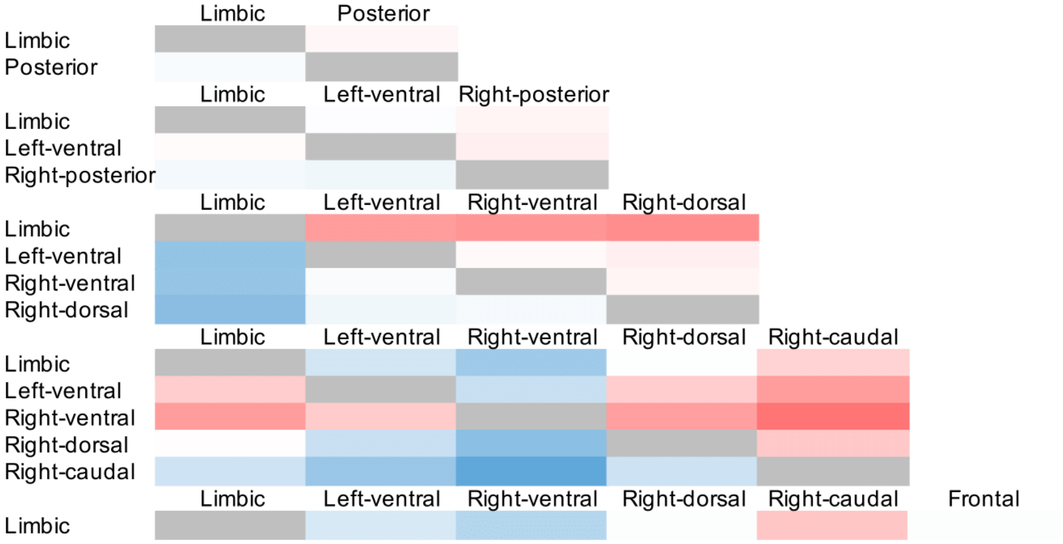

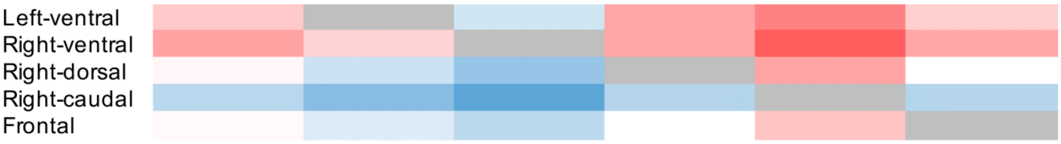


**Fragmented Letters**

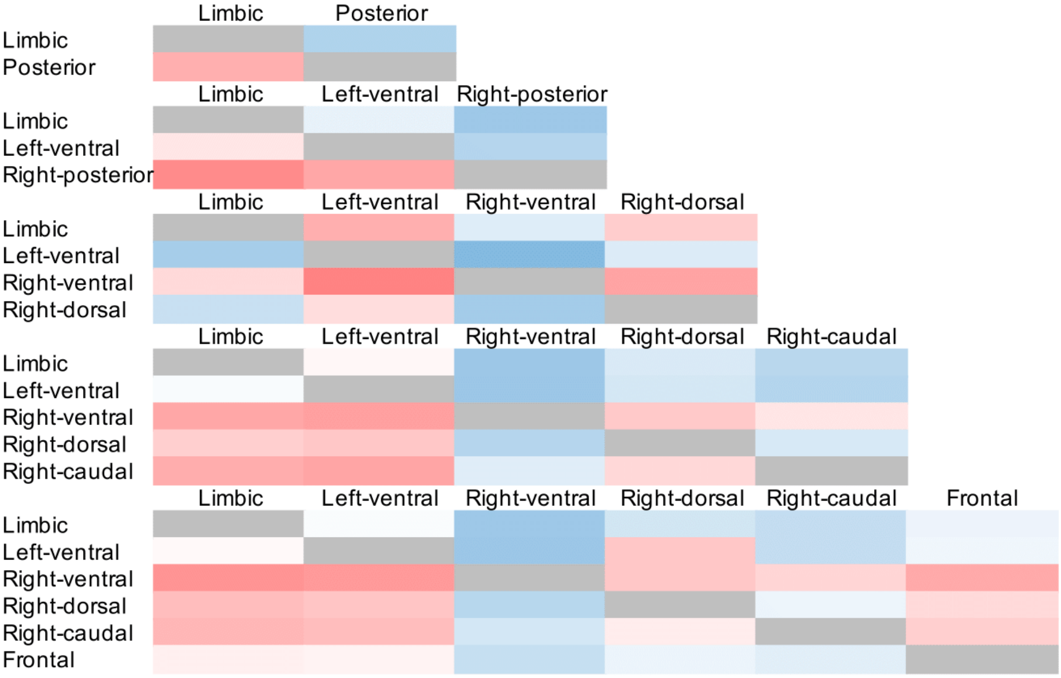


**Dot Counting**

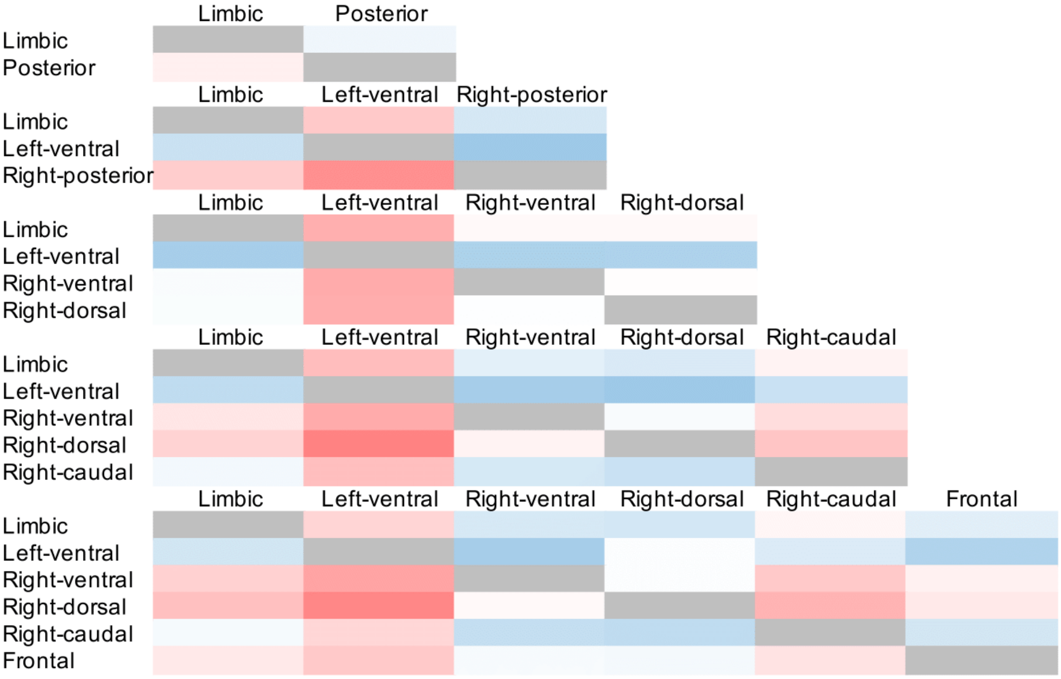


**Number Location**

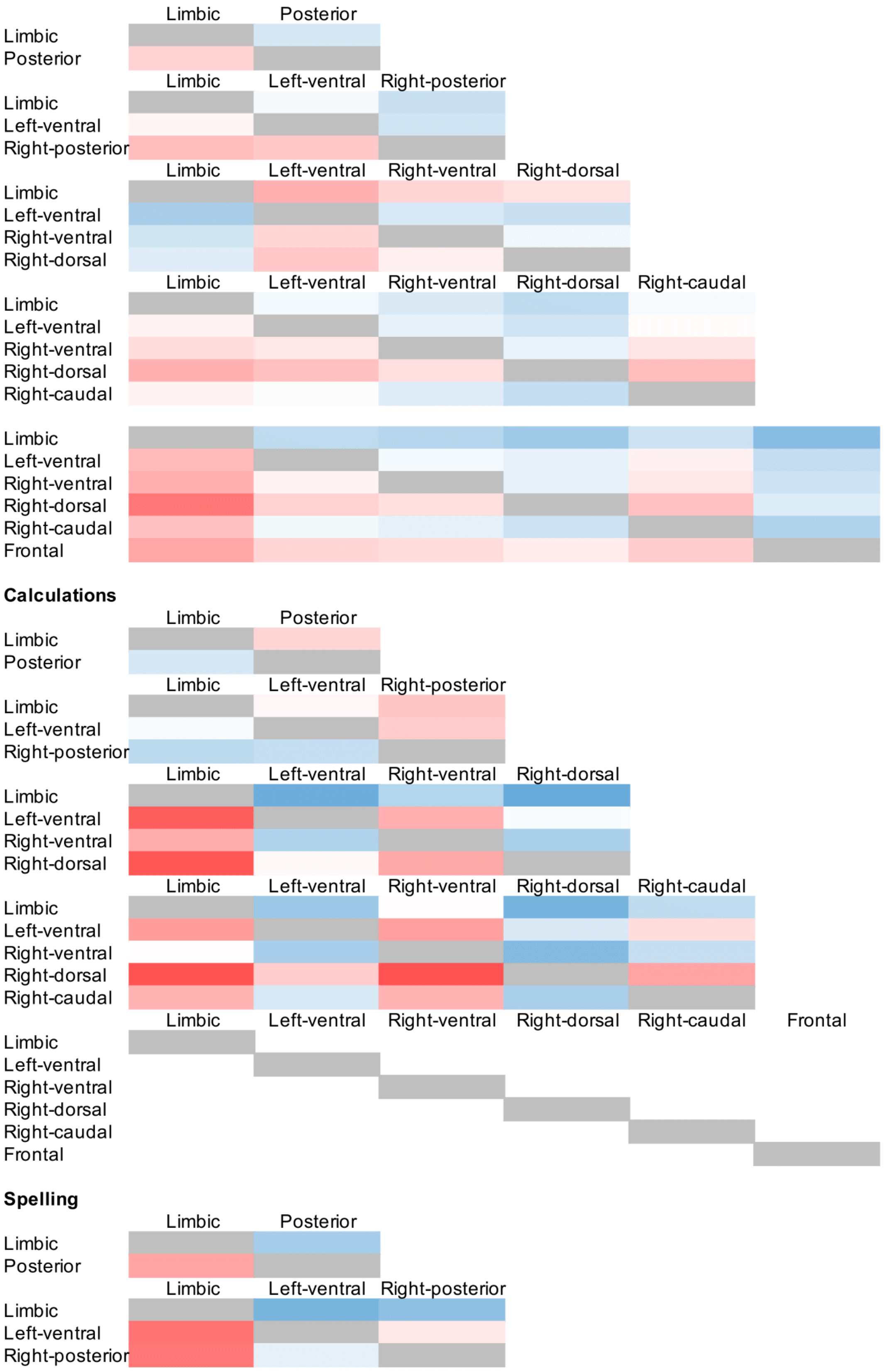
Limbic
Posterior
Left-ventral
Right-posterior
Right-ventral
Right-dorsal
Right-caudal
Frontal
Calculations
Spelling

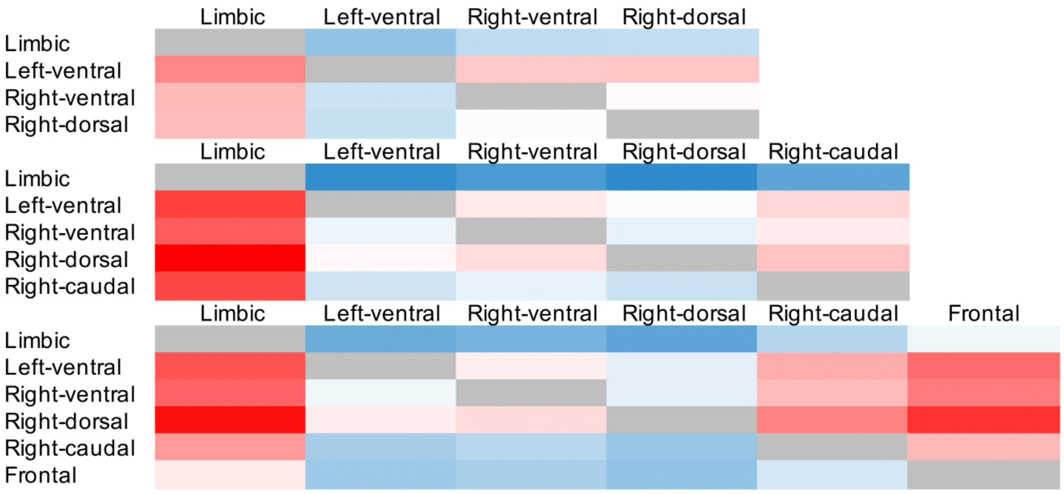


**Reading**

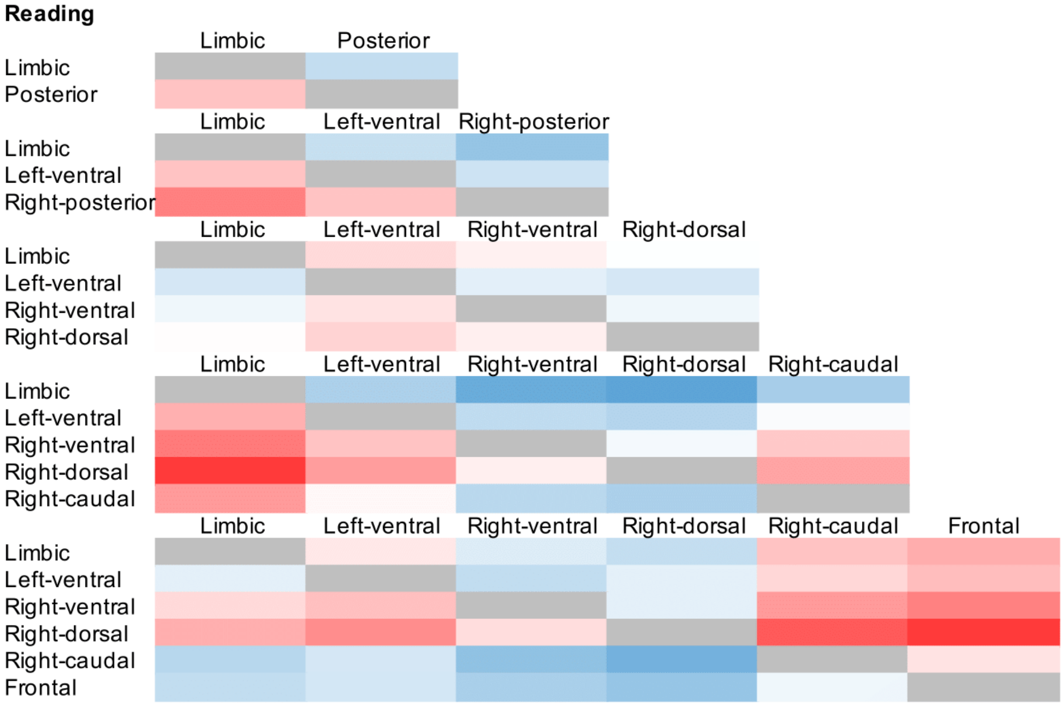


**Point Location**

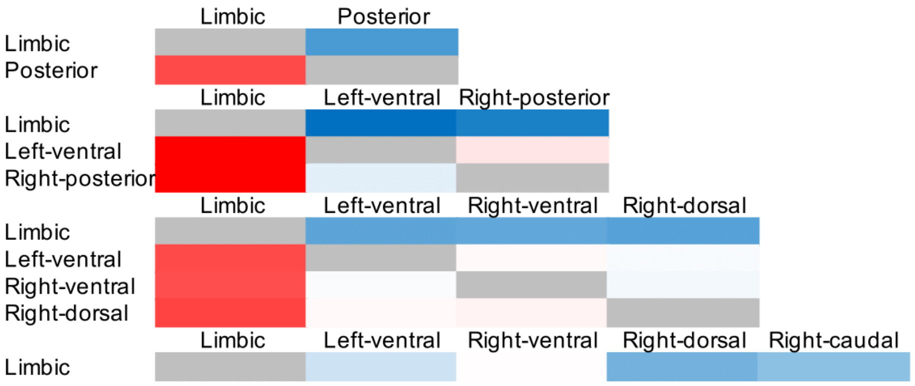

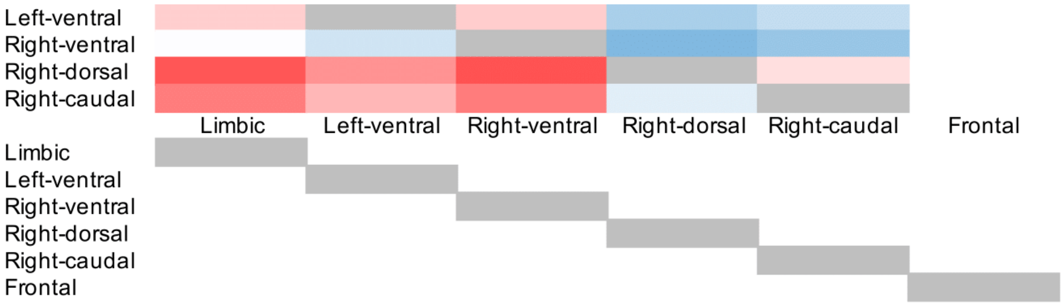


**Figure Discrimination**

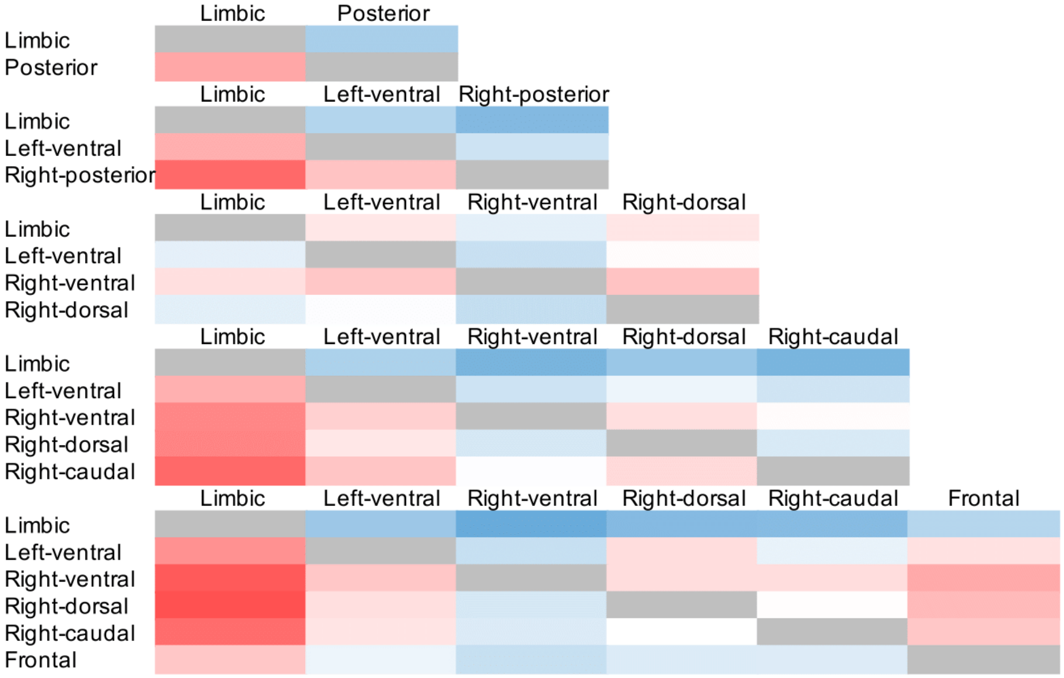


**Shape Discrimination**

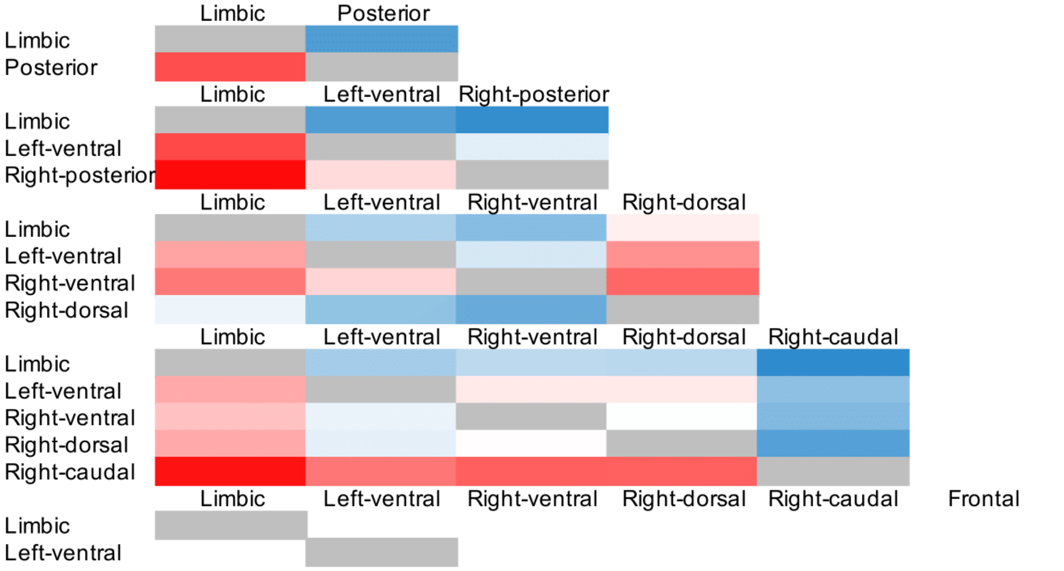

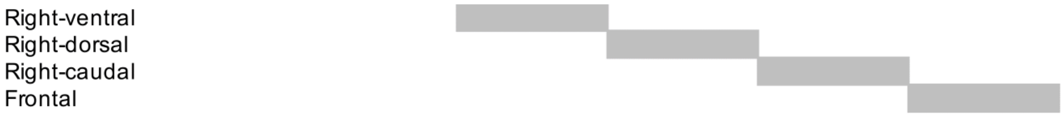


**Hue Discrimination**

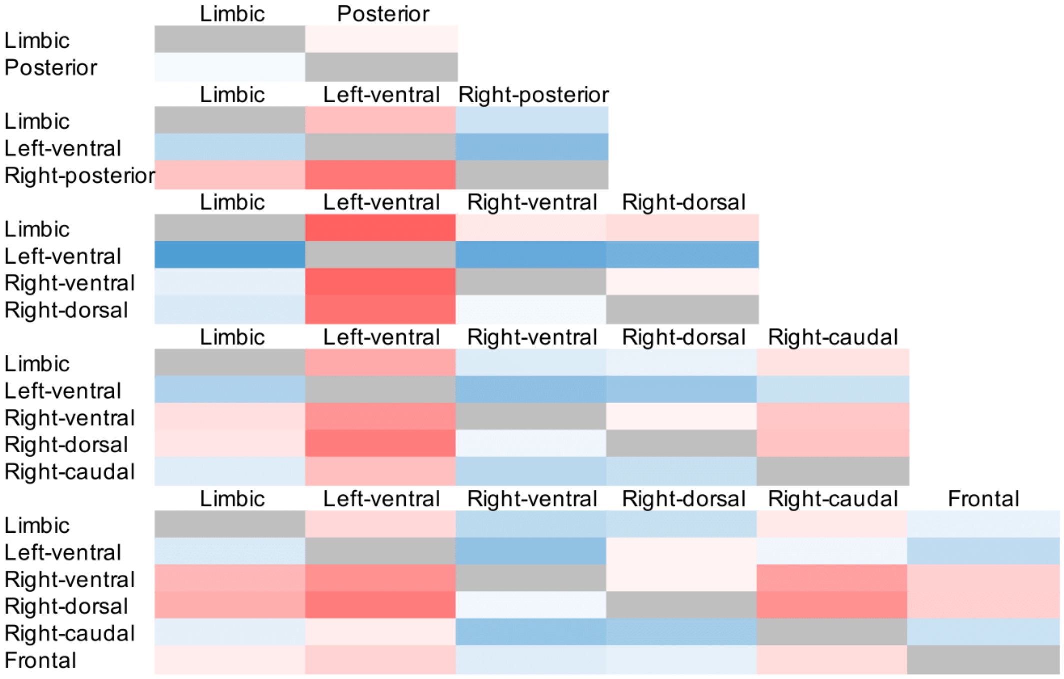


**Visual Acuity**

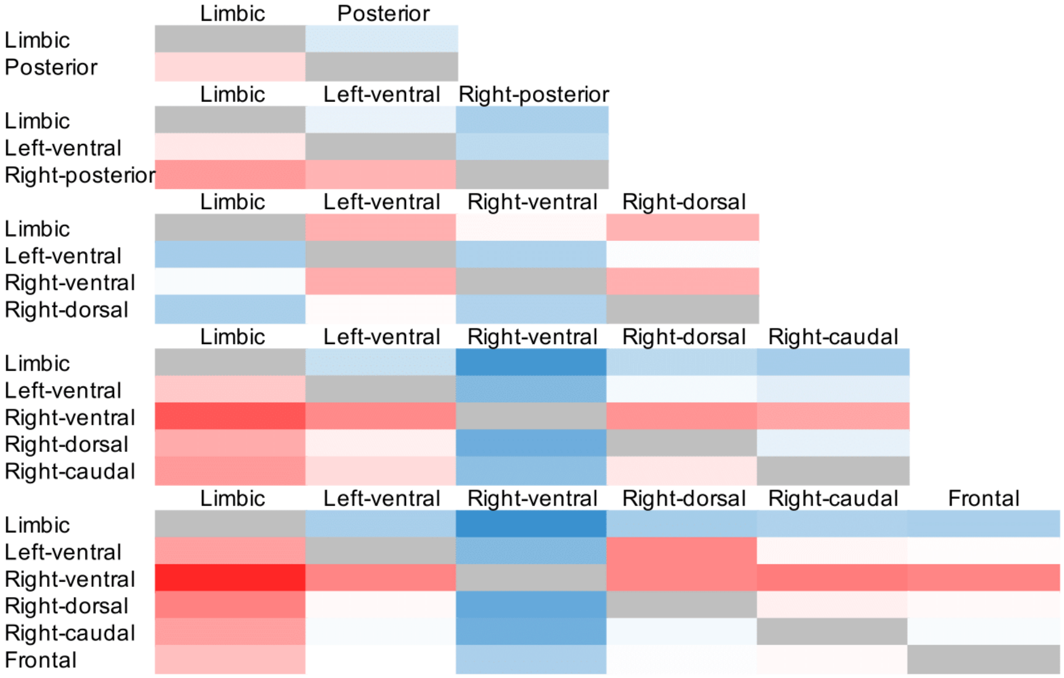


**Size Discrimination**

Limbic Posterior

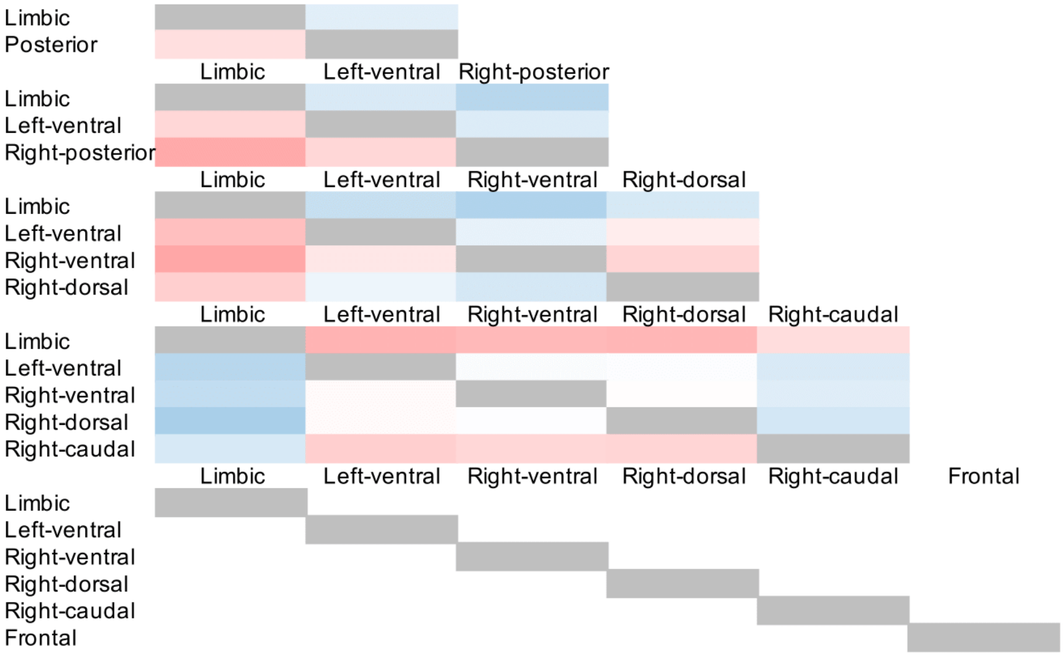


**Letter Cancellation**

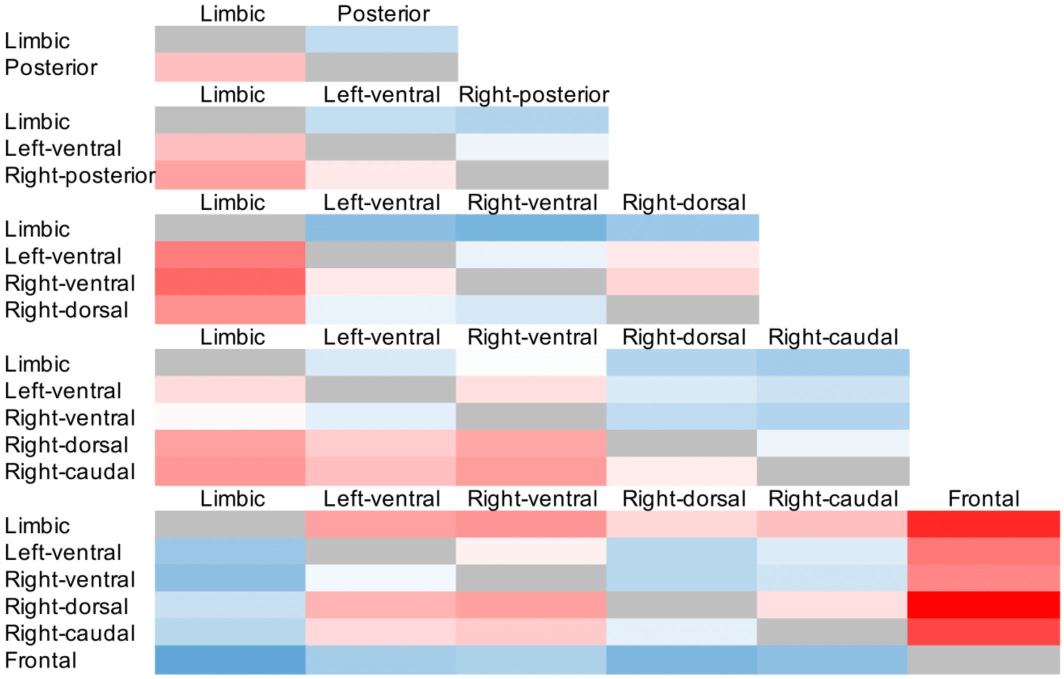


**Static Circle Detection**

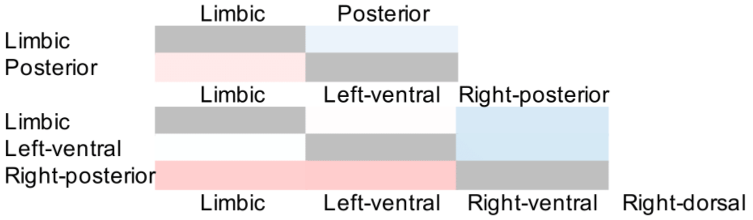

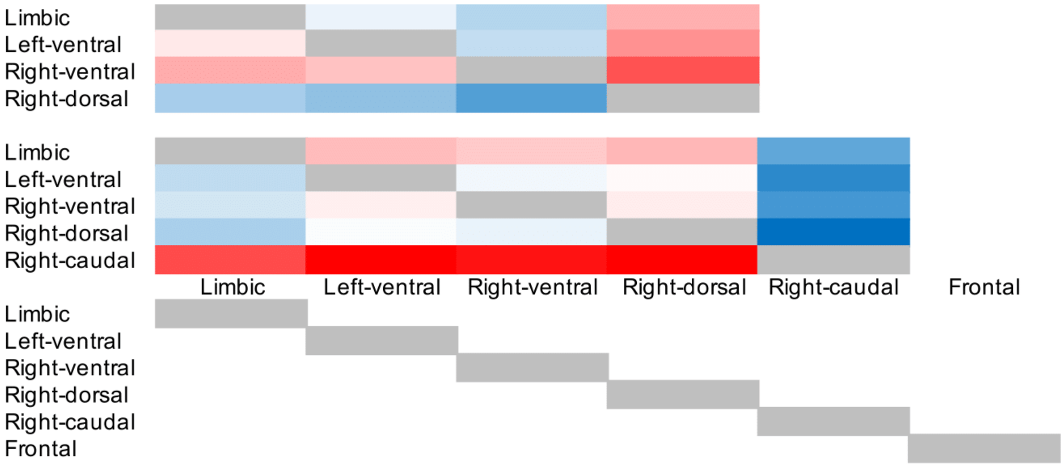


**Motion Coherence**

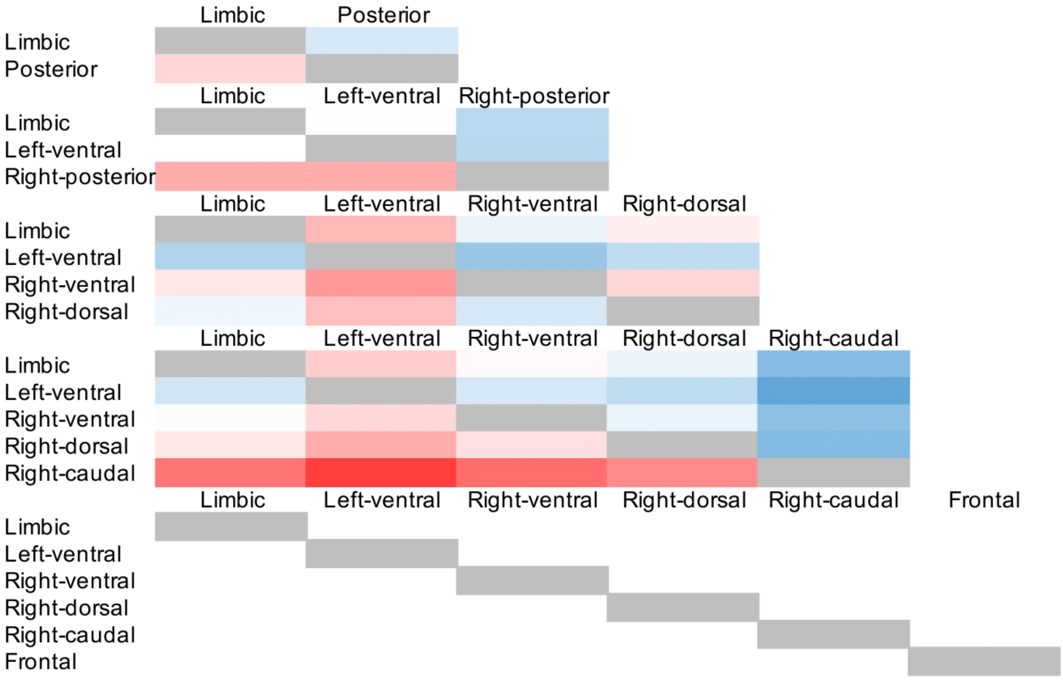

**Supplemental Figure 7. Associations between factor expressions from models *K*=2 through 6 and neuropsychological tests in the combined cohort**

The colors in the heatmap represent relative cross-sectional effects from the linear regression analyses. Red color indicates a negative association and blue a positive association. As in each model one of the factors is implicitly modelled. one of the factors serves as a reference (displayed horizontally) to assess effects of the factors displayed vertically. Empty cells indicate that the model could not be fitted due to insufficient observations and/or overfitting of the models.

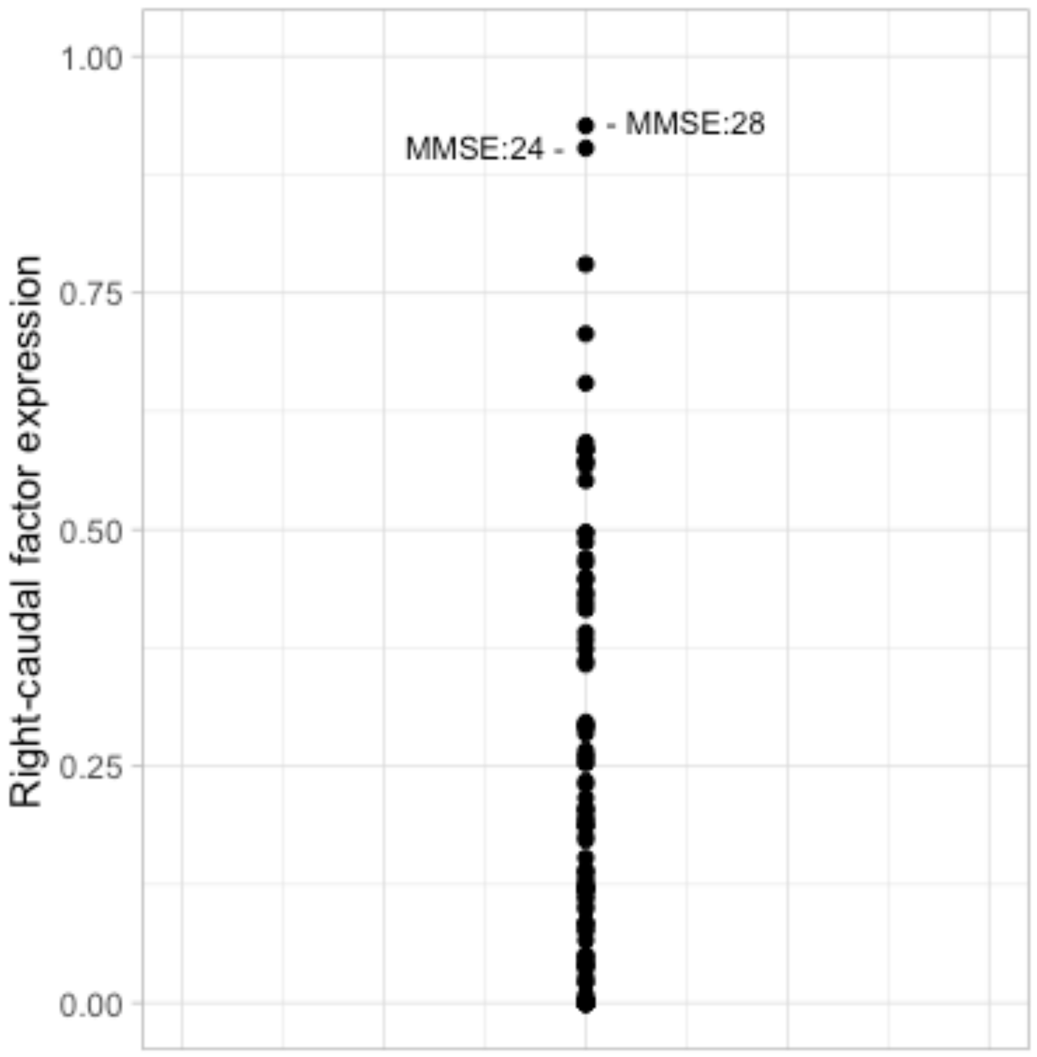

Right-caudal factor expression
1.00
0.75
0.50
0.25
0.00
MMSE:28
MMSE:24


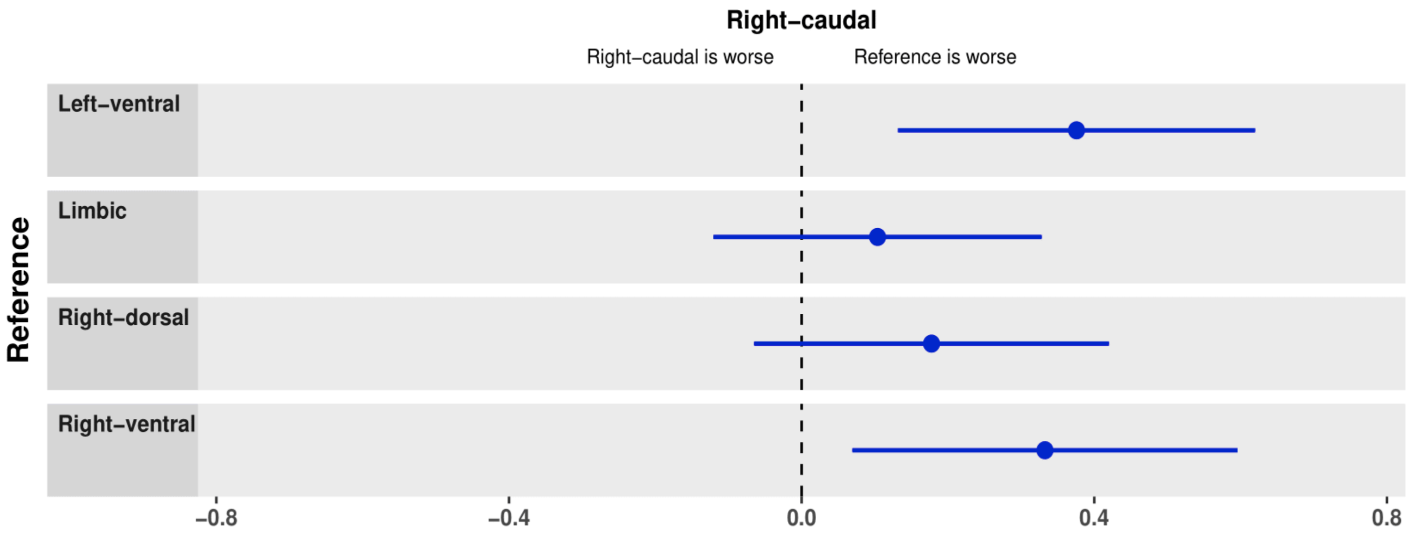

Right−caudal
Right−caudal is worse
Reference is worse
Reference
Left−ventral
Limbic
Right−dorsal
Right−ventral
−0.8
−0.4
0.0
0.4
0.8

**Supplemental Figure 8. Right-caudal factor observed in the *K* = 5 model**

Panel A displays the individual distribution of right-caudal factor expressions along the y-axis. The two outlies at the top were observed to have a high MMSE score, indicative being in an early clinical disease stage. Panel B contains relative cross-sectional effects from linear regression analyses. Lines indicate the 95% confidence intervals and a significant effect (uncorrected for multiple comparisons) is denoted by confidence intervals not including x=0. In these models the right-caudal factor is implicitly modelled and serves as a reference to assess effects of the others factors. Right-caudal factor expression is associated with better MMSE scores than the other factors, suggesting that high right-caudal factor expression might be a feature of early clinical disease severity in posterior cortical atrophy.

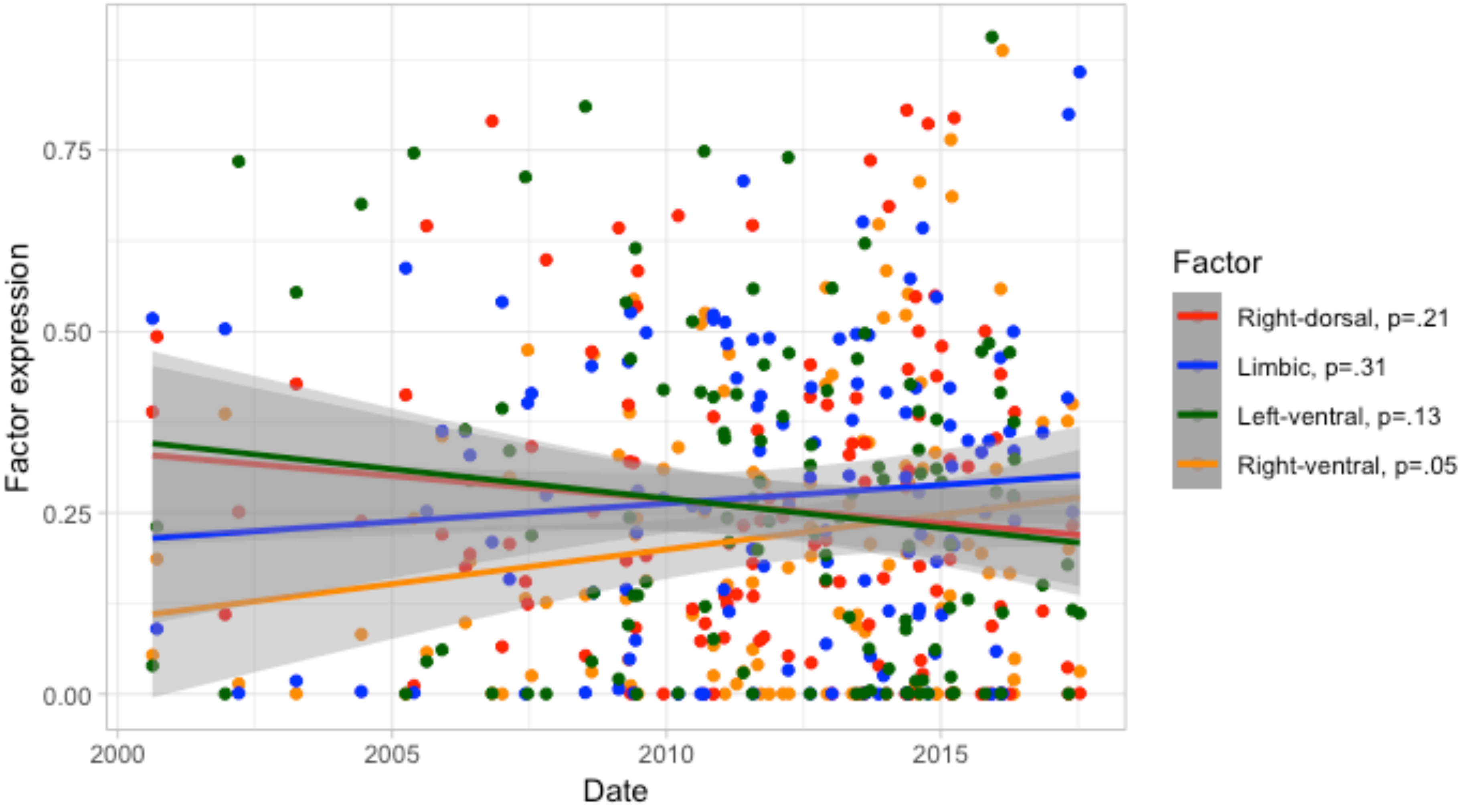

Factor expression
0.00
0.25
0.50
0.75
2000
2005
2010
2015
Date
Factor
Right-dorsal, p=.21
Limbic, p=.31
Left-ventral, p=.13
Right-ventral, p=.05

**Supplemental Figure 9. Factor expressions as a function of date of diagnosis**

This plot displays the date of diagnosis on the x-axis and factor expressions of the $K = 4$ model (combined sample) on the y-axis. None of the associations between factor expression and date were significant. This indicates that date of diagnosis, and in extension the clinical criteria used to determine diagnosis, was not related to the factor expressions obtained from the latent Dirichlet allocation models.

| | Right Dorsal vs Limbic | | | Right Dorsal vs Right Ventral | | | Right Dorsal vs Left Ventral | | |
|---|---|---|---|---|---|---|---|---|---|
| | β | SE | p | β | SE | p | β | SE | p |
| **Object perception** | | | | | | | | | |
| Fragmented Letters | 0.21 | 0.15 | 0.163 | 0.35 | 0.13 | **0.008** | -0.13 | 0.15 | 0.371 |
| **Space perception** | | | | | | | | | |
| Dot Counting | 0.03 | 0.16 | 0.865 | 0.01 | 0.14 | 0.965 | -0.32 | 0.15 | **0.031** |
| Number Location | 0.13 | 0.17 | 0.467 | -0.06 | 0.15 | 0.715 | -0.21 | 0.15 | 0.145 |
| **Non-visual dominant parietal** | | | | | | | | | |
| Calculations | -0.66 | 0.50 | 0.189 | -0.33 | 0.34 | 0.331 | -0.02 | 0.45 | 0.960 |
| Spelling | -0.26 | 0.43 | 0.544 | 0.01 | 0.29 | 0.964 | 0.22 | 0.47 | 0.641 |
| Reading | 0.00 | 0.42 | 0.994 | -0.06 | 0.28 | 0.835 | -0.17 | 0.53 | 0.755 |
| **Primary visual processing** | | | | | | | | | |
| Point Location | -0.74 | 0.70 | 0.293 | -0.05 | 0.70 | 0.948 | -0.02 | 1.03 | 0.982 |
| Figure Discrimination | 0.11 | 0.28 | 0.693 | 0.23 | 0.26 | 0.373 | 0.01 | 0.35 | 0.970 |
| Shape Discrimination | 0.07 | 0.38 | 0.856 | 0.59 | 0.27 | **0.027** | 0.43 | 0.40 | 0.287 |
| Hue Discrimination | 0.15 | 0.26 | 0.573 | 0.05 | 0.24 | 0.844 | -0.55 | 0.36 | 0.128 |
| Visual Acuity | 0.33 | 0.27 | 0.220 | 0.30 | 0.26 | 0.233 | -0.01 | 0.38 | 0.970 |
| Size Discrimination | -0.18 | 0.46 | 0.698 | 0.16 | 0.33 | 0.619 | 0.07 | 0.61 | 0.907 |
| Letter Cancellation | -0.43 | 0.26 | 0.093 | 0.16 | 0.24 | 0.492 | 0.08 | 0.32 | 0.800 |
| Static circle detection | 0.35 | 0.31 | 0.268 | 0.67 | 0.42 | 0.107 | 0.44 | 0.53 | 0.415 |
| Motion Coherence | 0.06 | 0.32 | 0.845 | 0.15 | 0.37 | 0.677 | -0.25 | 0.54 | 0.652 |
| **Memory** | | | | | | | | | |
| Verbal Learning Immediate | 0.26 | 0.13 | 0.051 | -0.01 | 0.13 | 0.939 | 0.01 | 0.12 | 0.936 |
| Verbal Learning Delayed | 0.43 | 0.12 | **0.001** | 0.16 | 0.12 | 0.179 | 0.23 | 0.11 | **0.036** |
| **Executive functions** | | | | | | | | | |
| Letter Fluency | 0.14 | 0.14 | 0.316 | -0.16 | 0.13 | 0.224 | -0.10 | 0.12 | 0.413 |
| Digit Span Forward | 0.10 | 0.14 | 0.474 | -0.24 | 0.14 | 0.084 | 0.16 | 0.13 | 0.210 |
| Digit Span Backward | 0.10 | 0.13 | 0.436 | 0.00 | 0.13 | 0.989 | 0.14 | 0.12 | 0.244 |
| **Language** | | | | | | | | | |
| Category Fluency | 0.46 | 0.13 | **0.000** | 0.04 | 0.12 | 0.763 | 0.06 | 0.11 | 0.610 |
| **Global cognition** | | | | | | | | | |
| MMSE | 0.32 | 0.12 | **0.007** | -0.02 | 0.12 | 0.880 | 0.03 | 0.11 | 0.746 |

| | Left Ventral vs Limbic | | | Left Ventral vs Right Ventral | | |
|---|---|---|---|---|---|---|
| | β | SE | p | β | SE | p |
| **Object perception** | | | | | | |
| Fragmented Letters | 0.34 | 0.17 | **0.043** | 0.48 | 0.14 | **<u>0.001</u>** |
| **Space perception** | | | | | | |
| Dot Counting | 0.34 | 0.17 | **0.044** | 0.32 | 0.15 | **0.030** |
| Number Location | 0.34 | 0.17 | 0.054 | 0.16 | 0.15 | 0.291 |
| **Non-visual dominant parietal** | | | | | | |
| Calculations | -0.63 | 0.53 | 0.231 | -0.31 | 0.47 | 0.516 |
| Spelling | -0.47 | 0.48 | 0.322 | -0.20 | 0.39 | 0.609 |
| Reading | 0.16 | 0.53 | 0.758 | 0.11 | 0.43 | 0.800 |
| **Primary visual processing** | | | | | | |
| Point Location | -0.70 | 0.88 | 0.425 | -0.02 | 0.63 | 0.974 |
| Figure Discrimination | 0.10 | 0.37 | 0.788 | 0.22 | 0.30 | 0.479 |
| Shape Discrimination | -0.36 | 0.42 | 0.391 | 0.16 | 0.35 | 0.641 |
| Hue Discrimination | 0.69 | 0.39 | 0.074 | 0.59 | 0.30 | **0.048** |
| Visual Acuity | 0.35 | 0.41 | 0.393 | 0.32 | 0.31 | 0.314 |
| Size Discrimination | -0.25 | 0.62 | 0.692 | 0.09 | 0.49 | 0.853 |
| Letter Cancellation | -0.51 | 0.34 | 0.135 | 0.08 | 0.28 | 0.767 |
| Static circle detection | -0.08 | 0.50 | 0.866 | 0.24 | 0.43 | 0.577 |
| Motion Coherence | 0.31 | 0.51 | 0.542 | 0.40 | 0.41 | 0.327 |
| **Memory** | | | | | | |
| Verbal Learning Immediate | 0.26 | 0.13 | 0.052 | -0.02 | 0.13 | 0.879 |
| Verbal Learning Delayed | 0.21 | 0.12 | 0.097 | -0.07 | 0.12 | 0.557 |
| **Executive functions** | | | | | | |
| Letter Fluency | 0.25 | 0.13 | 0.062 | -0.06 | 0.13 | 0.662 |
| Digit Span Forward | -0.06 | 0.15 | 0.680 | -0.41 | 0.14 | **0.004** |
| Digit Span Backward | -0.03 | 0.14 | 0.805 | -0.14 | 0.13 | 0.296 |
| **Language** | | | | | | |
| Category Fluency | 0.42 | 0.13 | **<u>0.001</u>** | -0.02 | 0.12 | 0.852 |
| **Global cognition** | | | | | | |
| MMSE | 0.30 | 0.12 | **0.011** | -0.05 | 0.12 | 0.643 |

| | Right Ventral vs Limbic | | |
|---|---|---|---|
| | β | SE | p |
| **Object perception** | | | |
| Fragmented Letters | -0.14 | 0.15 | 0.337 |
| **Space perception** | | | |
| Dot Counting | 0.02 | 0.15 | 0.888 |
| Number Location | 0.18 | 0.17 | 0.281 |
| **Non-visual dominant parietal** | | | |
| Calculations | -0.32 | 0.57 | 0.567 |
| Spelling | -0.27 | 0.43 | 0.527 |
| Reading | 0.06 | 0.42 | 0.896 |
| **Primary visual processing** | | | |
| Point Location | -0.69 | 0.82 | 0.400 |
| Figure Discrimination | -0.12 | 0.30 | 0.695 |
| Shape Discrimination | -0.52 | 0.37 | 0.163 |
| Hue Discrimination | 0.10 | 0.28 | 0.725 |
| Visual Acuity | 0.03 | 0.29 | 0.921 |
| Size Discrimination | -0.34 | 0.47 | 0.471 |
| Letter Cancellation | -0.59 | 0.27 | **0.028** |
| Static circle detection | -0.32 | 0.40 | 0.422 |
| Motion Coherence | -0.09 | 0.34 | 0.790 |
| **Memory** | | | |
| Verbal Learning Immediate | 0.26 | 0.13 | **0.037** |
| Verbal Learning Delayed | 0.26 | 0.12 | **0.027** |
| **Executive functions** | | | |
| Letter Fluency | 0.29 | 0.12 | **0.022** |
| Digit Span Forward | 0.32 | 0.13 | **0.012** |
| Digit Span Backward | 0.10 | 0.13 | 0.438 |
| **Language** | | | |
| Category Fluency | 0.41 | 0.12 | <u>**0.000**</u> |
| **Global cognition** | | | |
| MMSE | 0.33 | 0.11 | **0.003** |

**Supplemental Table 1. Tabular representation of cross-sectional and longitudinal associations between factor expressions and neuropsychological tests in the combined sample**

Values represent the relative cross-sectional associations between factor expression and neuropsychological tests. Negative values indicate that the factor under investigation is negatively associated with cognition compared to the reference factor in that comparison. while positive values indicate the opposite. Displayed p-values are bold when the effect is <0.05. uncorrected. P-values that remained significant after false discovery rate (FDR) correction are additionally underlined. FDR correction was applied according the number of tests and pairwise comparisons in this table (22*6=132).
